# Vertical inhibition of the autophagy pathway impairs growth and enhances sensitivity to mTORC1 inhibition in pancreatic ductal adenocarcinoma

**DOI:** 10.64898/2026.06.28.734981

**Authors:** Mallory K. Roach, Seamus E. Degan, Jonathan M. DeLiberty, Lily M. Pita, Noah L. Pieper, Runying Yang, Khalilah E. Taylor, Elyse G. Schechter, Ryan Robb, Mariaelena Pierobon, Clint A. Stalnecker, Emanuel F. Petricoin, Kirsten L. Bryant

**Author notes:** **Corresponding Author:** Kirsten L. Bryant: Lineberger Comprehensive Cancer Center, 450 West Drive, Chapel Hill, NC, 27599.

## Abstract

Pancreatic ductal adenocarcinoma (PDAC) is dependent on autophagy for growth. Chloroquine/Hydroxychloroquine (CQ/HCQ), the sole FDA-approved autophagy inhibitors, have shown limited clinical efficacy as cancer therapies. To identify approaches to improve PDAC response to CQ, we performed a CQ-anchored, CRISPR-Cas9 mediated loss-of-function screen. We identified that the loss of genes encoding proteins upstream in the autophagy pathway enhanced CQ-mediated growth suppression. This indicated that simultaneous targeting of two distinct nodes of the same pathway, vertical inhibition, may be a more effective strategy than single node inhibition. We demonstrated that genetic loss or pharmacological inhibition of VPS34, a protein necessary for autophagosome nucleation, sensitized PDAC cells to inhibitors of the terminal stage of the autophagy pathway, including CQ and an inhibitor of PIKfyve. We extended this concept to the initiation complex and demonstrated that ULK1/2 inhibition synergized with CQ and PIKfyve inhibition to impair PDAC cell growth and increase apoptosis. Anticipating mechanisms of resistance to vertical autophagy inhibition, we performed reverse-phase protein array profiling and identified that vertical inhibition of the autophagy pathway resulted in enhanced activation of the PI3K-AKT-mTORC1 signaling pathway. Increased mTORC1 signaling resulted in heightened sensitivity to bi-steric mTORC1 inhibition in both cell line and organoid models of PDAC. This study identifies novel anti-autophagy inhibitor combinations that may improve the clinical efficacy of autophagy inhibition for PDAC treatment.

**IMPLICATIONS:** Vertical inhibition of the autophagy pathway reduces pancreatic cancer cell growth, increases apoptosis, and enhances sensitivity to mTORC1 inhibition; thereby representing a novel therapeutic strategy for autophagy-driven pancreatic cancer.

## INTRODUCTION

Pancreatic cancer is characterized by a dismal 5-year survival rate of only 13% and limited therapeutic approaches (1–3). Pancreatic ductal adenocarcinoma (PDAC) accounts for over 90% of pancreatic cancer cases, making it the most common form of pancreatic cancer (4). PDAC is largely driven by mutations in the *KRAS* oncogene, which is critical for both tumor initiation and maintenance (5). Mutant KRAS orchestrates a variety of cellular signaling alterations that drive tumor growth, including metabolic reprogramming (6–8). In particular, KRAS mutant PDAC exhibits increased use of nutrient scavenging pathways, including macropinocytosis (9) and autophagy (10–12).

Autophagy is a multi-step catabolic process that leads to degradation and reuse of cellular components via trafficking to the lysosome (13). PDAC displays a heightened dependence on autophagy for both growth and survival (10,11). Consistent activation of the autophagy pathway generates critical bioenergetic intermediates that are necessary to maintain metabolic requirements and hence provide a proliferative advantage to pancreatic cancer cells in the nutrient-deprived PDAC tumor microenvironment (10). Furthermore, impairment of autophagy through both genetic suppression and pharmacological inhibition has been demonstrated to suppress growth in multiple models of PDAC (10,14). These preclinical findings led to the initiation of several clinical trials utilizing hydroxychloroquine/chloroquine (HCQ/CQ) (15,16), the sole FDA-approved inhibitors of autophagy, as monotherapy and in combination with standards of care for the treatment of PDAC (17–19). Although CQ/HCQ effectively inhibits autophagic flux in preclinical model systems, limited potency and efficacy have blunted their clinical therapeutic efficacy (17,20). Therefore, recent studies have focused on identifying combinatorial strategies with CQ to improve clinical outcomes (21,22). An alternative strategy is to identify more specific and/or potent mechanisms to inhibit the autophagy pathway in PDAC.

The autophagy pathway consists of four stages: initiation, nucleation, maturation, and degradation (13). Autophagy is initiated by the ULK1/2 serine/threonine kinase complex, which phosphorylates and activates other autophagy-related proteins in both the initiation and nucleation complexes, including ATG14 and VPS34 (13). The nucleation complex contains VPS34, a class III phosphoinositide 3-kinase (PI3K), which forms phosphatidylinositol 3-phosphate (PI3P) from phosphatidylinositol (PI). PI3P is a critical component of the forming the autophagosome (23). During the maturation stage, ATG7 and ATG3 process the scaffolding protein LC3B-I into mature LC3B-II, which is then integrated into the autophagosome membrane (13). Once the mature double-membrane autophagosome is formed, it then fuses with the lysosome, where the contents of the autophagosome are degraded for reuse (13). Pharmacological inhibitors of proteins within each stage of the autophagy pathway, including VPS34 (24) and ULK1/2 (25–28), have been developed, and ULK1/2 inhibitors have recently entered clinical trials (NCT04892017, NCT05957367). Additionally, we and others have demonstrated that inhibition of PIKfyve, a lipid kinase that is critical for autophagosomal and lysosomal fission and fusion, is an effective strategy to target the terminal stage of the autophagy pathway in PDAC (29,30).

Autophagy is regulated via the interplay of multiple nutrient-sensing kinases, including AMP-activated protein kinase (AMPK) and mammalian target of rapamycin complex 1 (mTORC1). AMPK senses cellular energy levels, and AMPK activation promotes autophagy during times of nutrient stress by phosphorylating inhibitory sites on mTORC1 and activating sites on ULK 1/2 (31). Conversely, in nutrient-replete conditions, active mTORC1 suppresses autophagy by phosphorylating negative regulatory sites on ULK1/2 and ATG13 (32). Due to its direct regulation of ULK1/2 phosphorylation, mTORC1 inhibition is a standard method for inducing autophagy (33).

Therapeutic targeting of mTOR signaling is of significant interest for the treatment of several cancers, including PDAC, as it is frequently activated and mediates several cancer-promoting pathways including metastasis and metabolism (34,35). While first generation mTORC1 specific inhibitors, known as rapalogs, have been approved for the treatment of some cancers, their clinical utility has been limited due to only partial inhibition of the key mTORC1 substrate 4EBP1 (32,36,37). Second generation mTOR inhibitors are ATP-competitive and bind both mTORC1 and mTORC2. These mTOR inhibitors more potently inhibit mTORC1 substrates; however, has limited their clinical efficacy (37,38). RMC-5552, a third-generation, bi-steric mTORC1 inhibitor, combines both allosteric and active state inhibitor modules from the first and second generation inhibitor classes, making it both potent and selective for mTORC1 (39,40). RMC-5552 has demonstrated tolerability and selective inhibition of mTORC1 in patients (NCT04774952) (41), with future studies aimed at identifying rational combinations for the treatment of cancer (41).

In the present study, we performed a CRISPR-Cas-9 genetic loss-of-function screen in the presence of CQ and identified that loss of multiple autophagy-related genes, including *PIK3C3*, the gene that encodes for VPS34, sensitized PDAC cells to CQ treatment. Based on this result, we hypothesized that concurrently inhibiting two nodes of the autophagy pathway, hereby referred to as “vertical autophagy inhibition,” may be a more efficacious strategy than single node inhibition. We found that pharmacological inhibition of VPS34 or ULK1/2 enhanced CQ- and PIKfyve inhibitor-mediated growth suppression. Growth suppression was due, in part, to increased apoptosis in combination treated cells. Vertical inhibition of the autophagy pathway also caused metabolic rewiring, resulting in enhanced PI3K-AKT-mTORC1 signaling. We demonstrated that this resistance mechanism could be therapeutically targeted with combined mTORC1 inhibition in both cell culture and organoid models of PDAC. We conclude that vertical inhibition of the autophagy pathway more effectively impairs autophagy in PDAC than single node inhibition, and that this combination strategy can be further enhanced by concurrent pharmacological inhibition of mTORC1. Ultimately, this therapeutic combination represents a novel anti-autophagy treatment strategy for PDAC.

## METHODS

### Cell Culture

Human KRAS-mutant cell lines Pa14C, Pa16C, Pa01C, and Pa02C generated from patient-derived xenograft (PDX) tumors, were supplied by A. Maitra (MD Anderson Cancer Center, Houston, TX). Each patient provided written informed consent, and studies were both authorized by an institutional review board and were carried out in compliance with one of the following recognized ethical guidelines: Declaration of Helsinki, CIOMS, Belmont Report, U.S. Common Rule. The following cell lines: HEK 293T (RRID: CVCL_0063), PANC-1 (RRID:CVCL_0480), MIA PaCa-2 (RRID:CVCL_HA89), and HPAC (RRID:CVCL_3517) were received from American Type Culture Collection (ATCC). All cell lines were cultured in DMEM (Gibco) containing 10% fetal bovine serum (FBS) and 1% penicillin-streptomycin solution. Cells were maintained in a 5% CO2 incubator at 37°C. Cell lines were not cultured past ten passages or one month and were frequently tested for mycoplasma using MycoAlert (Lonza). In 2017, all cell lines were profiled utilizing short tandem repeat profiling.

Patient-derived PDAC organoid models hT105 and hM1A were provided by Dr. David Tuveson (Cold Spring Harbor Laboratory) (Supplementary Table 1). The patient-derived PDAC organoid model PT3 was provided by Dr. Calvin Kuo (Stanford University) (Supplementary Table 1). PT3 organoid culture was established as previously described (42). After establishment, PT3 organoids were grown using the Matrigel dome method. Organoids were grown in a 5% CO2 incubator at 37°C. hT105, hM1A and PT3 organoids were grown in domes in growth factor reduced Matrigel (Corning) in complete human feeding medium consisting of: Advanced DMEM/F12 (Thermo Fisher Scientific) based WRN conditioned medium (L-WRN (ATCC CRL-3276)), 1x B27 supplement, 10 mM HEPES, 0.01 μM GlutaMAX (all from Thermo Fisher Scientific), 10 mM nicotinamide (Sigma-Aldrich), 50 ng/mL hEGF (Peprotech), 100 ng/mL hFGF10 (Peprotech), 0.01 μM hGastrin I (TOCRIS), 500 nM A83-01 (TOCRIS), and 1.25 mM (hM1A, hT105) or 1 mM (PT3) N-acetylcysteine (Sigma-Aldrich). PT3 organoid models were also grown in the presence of 10 μM SB202190 (Sigma), for the first two days of post-seeding PT3 organoid models were cultured in the presence of Y27632 (Selleckchem, 10.5 μM). For patient-derived cell lines and organoids, written informed consent was obtained from the patients. Patient studies were conducted in accordance with one of the Declaration of Helsinki. Patient studies were approved by an institutional review board.

### Antibodies, Plasmids and Chemical Reagents

The VPS34 inhibitor, SAR405 (S7682), PIKfyve inhibitor, apilimod (S6414), and ULK1/2 inhibitor, ULK-101 (S8793) were purchased from Selleckchem. Chloroquine diphosphate salt (CQ) (C6628) was purchased from Sigma-Aldrich. The mTORC1 inhibitor (RMC-5552) was purchased from MedChemExpress. Vinculin Antibody (V9264) was purchased from Sigma-Aldrich, VPS34 Polyclonal Antibody (GTX129528) was purchased from GeneTex. SQSTM1/p62 Antibody (5114), LC3B (2775), Phospho-Atg14 (Ser29) (92340), ATG14 (96752), 4EBP1 (9644), Phospho-4EBP1 (Thr37/46) (9459), S6 Ribosomal Protein (2317), Phospho-S6 Ribosomal Protein (S235/236)(2211) were purchased from Cell Signaling Technology, and all antibodies were used at the manufacturer’s suggested concentrations (Supplementary Table 1).

To generate *PIK3C3* knockdown PDAC cell lines, we utilized CRISPick from the Broad Institute to determine the most effective sgRNA sequence: sg-PIK3C3-A (AGTTGGGTTGGTGTAATGAG), sg-PIK3C3-B (TCGGTTGGTGCATCTAATGA), sg-PIK3C3-C (ATACACTCCCATATGGTGA). Oligos were amplified by PCR using Phusion^™^ Hot Start II DNA Polymerase (Thermo Fisher, F549S), forward primer (FWD: 5′-GGCTTTATATATCTTGTGGAAAGGACGAAACACCG-3′), and reverse primer (REV: 5′-CTAGCCTTATTTTAACTTGCTATTTCTAGCTCTAAAAC-3′). PCR-amplified oligos were assembled into the lentiCRISPRv2-Puro vector (Addgene, 52961) using the NEBuilder® HiFi DNA Assembly Master Mix (New England Biolabs, E2621). Two microliters of the assembly reaction were then transformed into DH5α competent cells (50 μl, Thermo Fisher Scientific, EC0112). From each transformation, four colonies were selected, and cultured. Plasmid DNA was isolated using the QIAGEN® Plasmid Plus Midi Kit (Qiagen, 12143). Plasmid DNA was then sequenced to ensure that desired sgRNA sequence was present. The resulting constructs were packaged into lentivirus as described below.

### Retroviral and lentiviral expression and vector infections

pBABE-puro-mCherry-EGFP-LC3B (Addgene plasmid #22418) and pMRX-IP-GFP-LC3-RFP-LC3ΔG (Addgene plasmid # 84572) stably expressing PDAC cell lines were generated as described previously (22,29). PDAC cell lines stably expressing the pMXs-puro-GFP-DFCP1 (Addgene plasmid #38269) were generated as follows: 900,000 HEK-293T cells were seeded in a T25 flask. Cells were then transfected the following day with a 1:3 ratio of plasmid DNA to Fugene6 (Promega, #E2691), 1.67 µg of packaging vector gag/pol (Supplementary Table 1) and 0.835 µg of packaging vector VSGg (Supplementary Table 1) mixed in 400 µL of OptiMEM (Gibco). HEK-293T cells incubated with DNA/Fugene transfection mixture overnight and then media was changed to DMEM with 20% FBS and retroviral generation proceeded for 48 hours. 48 hours post media change viral supernatant was harvested and filtered with a 0.45-micron filter and immediately was added to PDAC cell lines in T-25 flasks that were 30-50% confluent on the day of transduction. Polybrene was added to the filtered virus solution so that the final concentration of polybrene in the flask was 8 µg/mL. Flasks were incubated with retrovirus for 8 hours and then retrovirus was diluted with 5 mL of media. Once T25 flasks reached confluency, cells were selected using 1-5 µg/mL of puromycin preselected based on cell line kill curves. Cells were then trypsinized and moved to 10cm plates and were then selected until no cells remained adhered in kill plate.

For lentiviral packaging of CRISPR knockdown guides, HEK-293T cells were seeded in T25 flasks as described above. Cells were then transfected the following day with a 1:3 ratio of plasmid DNA to Fugene6 (Promega #E2691), 3 µg of packaging vector psPAX2 (Supplementary Table 1) and 1 µg of packaging vector pMD2.G (Supplementary Table 1). These were mixed in 400uL of OptiMEM (Gibco). After being incubated at room temperature for 15 minutes, this mixture was added dropwise onto cells. Following 18 hours of incubation, media was changed to 5.5 mL of DMEM (Gibco) with 20% FBS. Viral supernatant was harvested 48 hours later and aliquoted and store at -80° C until it was used. PDAC cells were seeded to 80% confluence and virus was added with 8 µg/mL polybrene in media. Flasks were incubated with lentivirus for 8 hours and then lentivirus was diluted with 5 mL of media. Once T25 flasks reached confluency, cells were selected using 1-5 µg/mL of puromycin as described above. Following approximately 7 days of selection, cells were seeded for proliferation and western blotting assays.

### CRISPR-Cas9 screen

A CRISPR-Cas9 loss-of-function screen was performed by infecting four PDAC cell lines (Pa01C, Pa02C, Pa14C, PANC-1) with a barcoded druggable genome CRISPR library as previously described (43,44). Briefly, cells were infected with the druggable genome library at an MOI of 0.2 and cells were selected using puromycin. Genomic DNA was harvested on day 0 (2 days post-infection) in order to elucidate coverage of the library. Genomic DNA was also collected on day 7 (following stable guide incorporation). After guides were stably incorporated, cells were split into two treatment groups vehicle and 3.125 µM of CQ. After four weeks of treatment (day 35), cells were harvested, and genomic DNA (gDNA) was isolated using a DNeasy Blood & Tissue Kit (Qiagen, 69504). From the gDNA, sgRNA was amplified and analyzed via next generation sequencing. sgRNA abundances were quantified, and gene essentialities were determined using MAGeCK analysis (45). Sample quality was evaluated via principal component analysis (PCA) and Spearman correlations of median normalized sgRNA abundances. The screen was further validated by assessing the genetic dependencies of established essential and non-essential genes.

### Immunoblotting

Phosphatase Inhibitor Cocktail Set I (Millipore; 524624) and Set II (Millipore; 524625) and Protease Inhibitor Cocktail (Roche; 11873580001) were freshly added to 1% Triton buffer (25 mM Tris-HCl, pH 7.4, 100 mM NaCl, 1 mM EDTA, 1% Triton) to make Lysis buffer. On ice, Cells were washed twice with 1X PBS then lysed using Lysis buffer. Cells were then scraped and collected in prechilled Eppendorf tubes and centrifuged at 14,000 g for 10 minutes at 4° C to generate clarified lysates. Lysates were then quantified and standardized to an equal concentration using Pierce BCA Protein Assay Kit (Thermo Fisher 23227). Standardized proteins were separated using SDS-PAGE electrophoresis and transferred to PVDF membranes. Membranes were blocked at room temperature in 5% (w/v) BSA in TBS with 0.05% Tween 20 (TBST) for 1 hour. Following 1 hour blocking, membranes were placed in primary antibody and incubated rocking overnight at 4° C. The next day, membranes were washed in TBST three times for ten minutes each and then incubated with secondary antibody for 1 hour at room temperature. After 1 hour in secondary antibody, membranes were washed again with TBST three times and protein levels were determined by chemiluminescence.

### Fluorescent microscopy

PDAC cell lines expressing the GFP-DFCP1 probe were seeded in a 35 mm glass bottom dish and allowed to adhere overnight. Cells were then treated for desired time points with inhibitor or control. Prior to imaging, cells were incubated with Hoechst (1:2000) to allow for nuclei labelling. Cells were imaged live on the Zeiss 900 confocal microscope at 40X magnification and excited with 488-nm laser lines. After 10 images were acquired, results were analyzed in FIJI. To determine the GFP-DFCP1 positive area per cell, the total fluorescence of GFP was quantified in FIJI following the removal of background fluorescence via thresholding. The number of nuclei per image were then counted in Fiji and the ratio of GFP area per cell number was determined for each image. The average of all ten images was then normalized to the average for the DMSO control for each biological replicate.

PDAC cell lines expressing the mCherry-EGFP-LC3B probe were seeded in a 35 mm glass bottom dish and allowed to adhere overnight. Cells were then treated for desired time points with inhibitor or control. Cells were imaged live on the Zeiss 900 confocal microscope at 40X magnification and excited with 488- and 561-nm laser lines. After 10 images were acquired, results were analyzed in FIJI as previously described (22). Briefly, following thresholding to capture labeled puncta, the total area of mCherry positive fluorescence puncta and EGFP positive fluorescent puncta per image was determined in FIJI and the ratio of mCherry fluorescent area to EGFP fluorescent area was used to determine the level of autophagic flux. The average of all ten images was then normalized to the average for the DMSO control for each biological replicate.

### Flow cytometry

PDAC cells stably expressing GFP-LC3-RFP-LC3ΔG probe were plated in 6-well plates and treated with the specified treatment conditions. Following the desired drug incubation time, washed twice with PBS, lifted in trypsin and collected in media in 1.7mL Eppendorf tubes. Samples were centrifuged at 500 g at room temperature for 5 minutes. The supernatant was removed, and the cell precipitate was resuspended in 500-600 µL of media and passed through a cell sorting cap into a flow cytometry tube. Samples were analyzed using an Attune Flow Cytometer from Thermo Fisher Scientific to quantify total GFP and RFP fluorescence. Autophagic index is represented as a ratio of RFP/GFP.

Apoptosis assays were performed in PDAC cell lines using the Invitrogen Dead Cell Apoptosis Kit with Annexin V FITC & Propidium Iodide (V13242) according to manufacturer’s instructions. Briefly, media from each well of a 6-well plate was collected and transferred to 10 mL conical tubes (Corning). 500 µL of PBS was added to each well, plate was swiveled, and PBS was collected and placed into the 10 mL conical tube containing collected media. 250 µL of benchtop trypLE was added to each well, and the plate was incubated for 15 minutes at 37°C. After incubation, the previously collected media (∼500 µL) was used to collect the trypsinized cells. Cells were then pelted by centrifugation at ∼500 × g for 5 minutes at room temperature. The media was aspirated carefully, ensuring the pellet remained undisturbed. 1 mL of PBS was added to each 10 mL Falcon tube to resuspend the cell pellet, which was then transferred into a 1.7 mL Eppendorf tube. Tubes were centrifuged again at 500 × g for 5 minutes at room temperature. The PBS was carefully aspirated without disturbing the pellet and each sample was then resuspended in 25 µL of Annexin V–PI staining solution containing 10x binding buffer (1:10 dilution), propidium iodide (1:1000 dilution), Annexin V–FITC (1:200 dilution) in ddH₂O. Samples were vortexed gently and incubated in the dark for ∼15 minutes. After incubation, 400 µL of the 1× binding buffer was added to each sample and mixed well.

Samples were processed on a CytoFLEX flow cytometer within one hour to ensure optimal signal detection. Cytobank was used to analyze the acquired FCS files as previously described (29). Briefly, FCS files were uploaded to Cytobank, and cells were gated to exclude debris and doublets. Then additional gating was optimized using the DMSO condition of each cell line to create four quadrants for PI-area and FITC-area. This gating was applied to the treatment conditions, the resulting sum of the lower right and upper right quadrant (FITC-high, PI-low and FITC-high, PI high) was used to determine the percentage of early and late apoptosis.

### Drug response testing

2-D proliferation assays were performed by seeding 750-1000 cells (depending on cell line) in a 96-well overnight. The cells were drugged at the desired concentration 24 hours after seeding. After 5 days, Hoechst (nuclei stain) was added to the cells and incubated for 15-30 minutes. Cells were counted using a BioTek Cytation 1 Cell Imaging Multi-Mode Reader. All cell counts were normalized to control DMSO cell count as 100% and the cell count at day 0 (the day of drugging) for 0%.

Long-term colony formation growth assays were performed by seeding 1,000 cells in a 6-well plate. The following day, cells were drugged at desired concentration. Every 3-4 days media was refreshed containing the desired drug concentrations. Cells were fixed following 10-14 days of treatment, depending on when the vehicle treated condition reached confluency. First, cells were washed in 1X PBS and then cells were fixed by adding a crystal violet fixation solution made up of 1g of crystal violet per 100mL of 1X PBS with 4% paraformaldehyde. After cells incubated with fixing solution for 30 minutes, plates were washed gently with water and dried for 2 days prior visualization with an Alexa Flour 647 filter (Typhoon FLA 9500). Fluorescent images were then analyzed in FIJI by thresholding for the crystal violet fluorescence and calculating the percent area covered by crystal violet for each well. Results were normalized to DMSO control as 100%.

For drug response testing in 3D organoid models of PDAC, a suspension of approximately 1000-2000 organoids per/well was plated in a 96 well plate with white wells. Organoids were plated in 150 µL of DMEM with 10% Matrigel and 1:1000 Y-compound and were seeded for 48 hours prior to drugging. Organoids were then drugged for 5 days, and viability was determined via the CellTiter-Glo 3D Viability Assay (G9682) from Promega according to the manufacturers protocol.

### Sample preparation and reverse phase protein array (RPPA)

PDAC cells were plated in six-well plates and treated with DMSO, CQ (6.25 µM), apilimod (100 nM), ULK101 (3 µM), concurrent CQ and ULK101 (6.25 µM and 3 µM respectively), or concurrent apilimod and ULK101 (100 nM and 3 µM respectively) for 4, 24 or 72 hours. Samples were lysed and RPPA was performed as previously described (22,29,46–48) Principal component analysis (PCA) was used to determine how replicates, cell lines, treatments, and timepoints clustered. No samples were excluded from the final analysis. Antibodies that did not have determined intensities or had intensities less than one were removed on a sample-by-sample basis from the final analyses. Intensities were Log_2_ transformed and then median normalized. Limma (v.3.56.2) (Supplementary Table 1) analysis was performed for differential expression analysis in R-studio (V.4.4.2) (49). The “all” cell line condition was created with individual cell lines used as blocking terms and all plots were generated in R-studio (50)

### Statistical analysis

All data was analyzed by GraphPad Prism (Supplementary Table 1) and is normalized to each dataset’s respective controls. All data was replicated with n≥3 independent experiments. For all graphs, error bars are representative of mean or median +/- SEM or SD. Where significance is notated, one asterisk (*) indicates a significance p < 0.05, two asterisks (**) indicates a significance p < 0.01, three asterisks (***) indicates a significance p < 0.001 and four asterisks (****) indicates a significance p < 0.0001. The number of samples analyzed for each experiment and whether the data presented is an average of multiple technical replicates and/or normalized is clarified in the figure legend for each experiment.

### Data availability

Data from this research that are not available within this paper, including the processed reverse phase protein array (RPPA) data and CRISPR-Cas9-loss-of-function screen data, are available upon request from the corresponding author.

## RESULTS

### CRISPR-Cas9 loss-of-function screen identifies genes that modulate CQ sensitivity

To identify therapeutic targets with the potential to enhance CQ-mediated growth suppression in PDAC, we performed a pooled, lentiviral CRISPR-Cas9 mediated loss-of-function screen using a library of 2,240 genes related to cancer signaling pathways (43). Following library transduction, four *KRAS*-mutant PDAC cell lines were cultured for 28 days in the presence of a sublethal dose of CQ (3.125 μM) or vehicle (**Fig. 1A**). Samples were sequenced, and the relative enrichment and depletion of sgRNAs were evaluated (**Fig. 1B**). Principal component and Spearman correlation analyses were utilized to confirm sample quality (Supplementary Fig. S1A and B). MAGeCK analysis was used to measure changes in sgRNA abundance for all genes (45). To validate the screen, we compared the MAGeCK β-scores of both essential and non-essential genes for the vehicle treated control cells, as previously described (21), and found that essential genes were consistently lost in control cells, while genes predicted to be non-essential were less frequently lost (Supplementary Fig. S1C).

**Figure 1.**
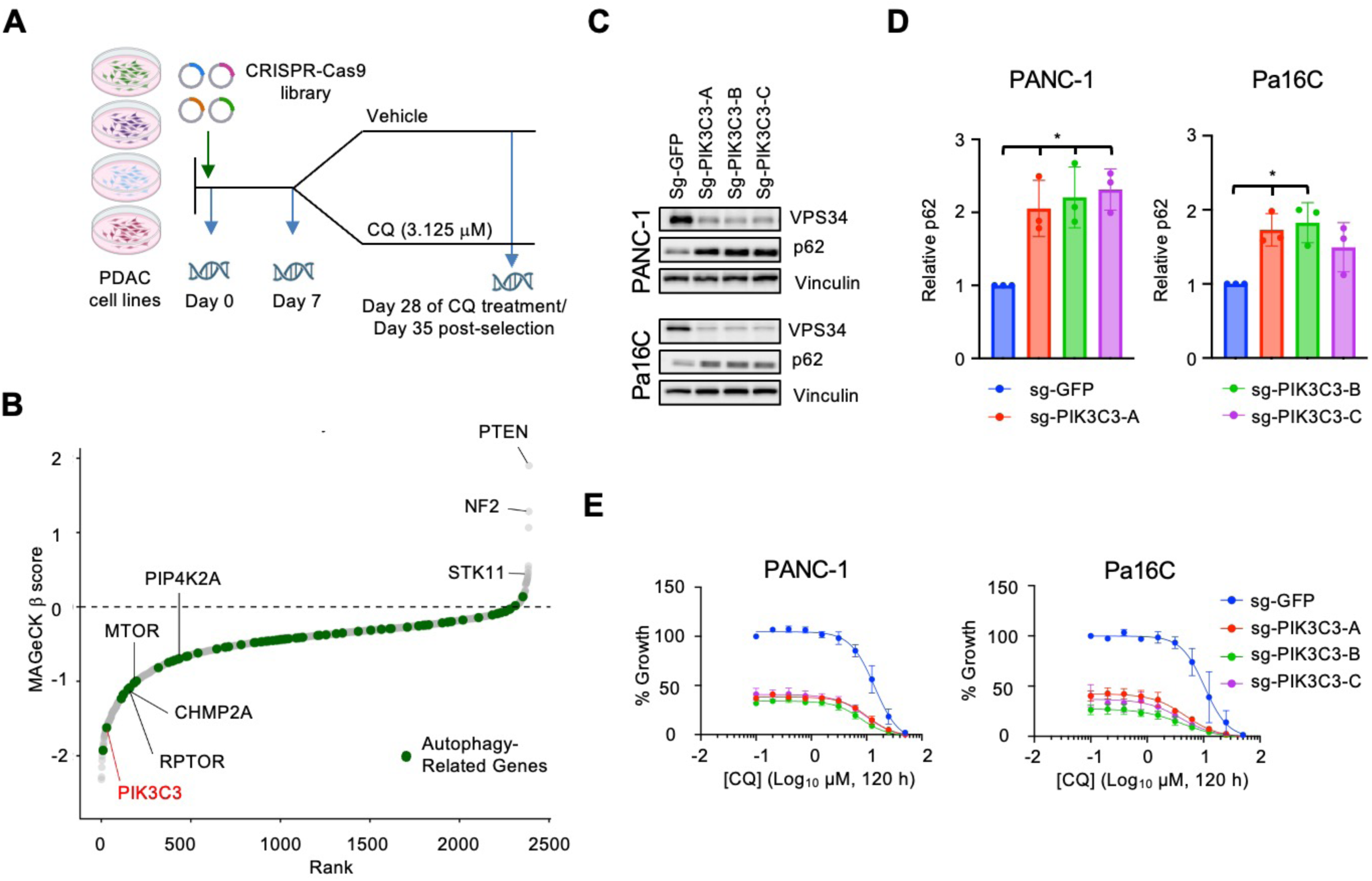
Genetic suppression of *PIK3C3* sensitizes *KRAS*-mutant PDAC cells to CQ treatment. **A,** Schematic of CRISPR/Cas9 loss-of-function screen. **B,** CRISPR/Cas9 loss-of-function screen performed in PANC-1, Pa01C, Pa14C and Pa02C cells treated with CQ (3.125 µM) for 28 days. Autophagy-related genes are in green. Genes are ranked by relative enrichment/depletion (β-scores) in the CQ condition. **C,** Immunoblot of PANC-1 and Pa16C cell lines following *PIK3C3* knockdown. Blots are representative of three independent experiments. **D,** Densitometry of relative p62/SQSTM1 protein levels from blots in **C** normalized to sg-GFP control. Error bars represent the mean ± SD of three independent experiments. Statistical significance was determined by a one-way Student’s *t-test*. *, π < 0.05. **E**, Five-day viability assay in PANC-1 and Pa16C cells infected with control guide (sg-GFP) or three distinct sgRNAs against *PIK3C3* (sg-PIK3C3-A, sg-PIK3C3-B, sg-PIK3C3-C) and treated with a dose response of chloroquine [0.2 – 50 µM]. Each data point represents mean ± SEM of three independent experiments.

Next, we evaluated sgRNAs enriched in our CQ-treated cells to identify genes that could confer resistance to CQ treatment. We found enrichment of several tumor suppressor genes previously demonstrated to confer resistance to targeted therapies in PDAC, including *PTEN*, *NF2*, and *STK11* (**Fig. 1B**). PTEN negatively regulates the PI3K-AKT-mTOR signaling pathway, which is a critical signaling pathway in PDAC that can be directly activated by mutant KRAS to promote cell survival and tumor growth (35) and directly regulates autophagy (51). Additionally, *PTEN* loss is associated with PDAC tumor development and increased NF-κB signaling, which leads to activation of cytokines and enhances the inflammatory tumor microenvironment (52). *STK11/*LKB1 acts upstream of AMPK to regulate metabolism in response to nutrient levels and *STK11/*LKB1 loss facilitates tumor growth by decoupling nutrient sensing from activation of metabolic pathways (53) and increases metastasis in PDAC (54). Loss of *STK11* or *PTEN* or conversely activating mutations in the LKB1 or PI3K pathways may represent biomarkers of resistance to anti-autophagy therapies.

To identify potential combination therapies, we next evaluated genes for which sgRNAs were depleted rather than enriched. Inhibition of the proteins these genes encode would sensitize PDAC cells to CQ treatment. The library contained sgRNAs to 72 autophagy-related genes. Interestingly, most of these genes were significantly depleted in CQ-treated cells (**Fig. 1B**; Supplementary Table 2). Among the genes with depleted sgRNAs, we found multiple components of the mTOR pathway (*MTOR, RPTOR* and *IGF1R*) (**Fig. 1B**; Supplementary Table 2) which are well established regulators of autophagy (55), and have previously been identified as sensitizers to CQ treatment in PDAC (21). This validated our screen to identify modulators of the autophagy pathway that enhance sensitivity to CQ.

Among the most significant hits from the screen was *PIK3C3* (**Fig. 1B**). *PIK3C3* encodes VPS34, which associates with VPS15 and ATG14L to form VPS34 Complex I, a critical mediator of autophagosome nucleation (23,56). To validate that loss of *PIK3C3* sensitized PDAC cells to CQ-mediated growth suppression, we generated two PDAC cell lines that stably expressed three distinct sgRNA’s targeting *PIK3C3.* To confirm that genetic suppression of *PIK3C3* impaired autophagy, we immunoblotted for p62/*SQSTM1*, a chaperone protein that associates with protein aggregates and facilitates their targeted degradation via the autophagic pathway (57). Accumulation of p62/*SQSTM1* is a canonical marker of autophagy suppression (58). We found that genetic suppression of *PIK3C3* led to the accumulation of p62/*SQSTM1* (**Fig. 1C** and **D**) indicative of impaired autophagy. Furthermore, we observed that genetic suppression of *PIK3C3* led to the accumulation of cellular vacuoles which are canonically associated with VPS34 loss and autophagy inhibition (59,60) (Supplementary Fig. S1D). Importantly, we also observed that stable suppression of *PIK3C3* resulted in decreased proliferation, and this was further potentiated by the addition of CQ (**Fig. 1E**). In summary, genetic suppression of *PIK3C3* impaired autophagy and sensitized PDAC cells to CQ treatment.

This finding led us to hypothesize that vertical inhibition of the autophagy pathway may more strongly decrease PDAC cellular proliferation relative to single node inhibition. While this hypothesis has not been tested in the context of autophagy inhibition, vertical inhibition of a signaling cascade has been well documented as a potential therapeutic strategy for targeting the RAS ERK-MAPK pathway across multiple RAS-driven cancers (61,62), including pancreatic cancer (43). In fact, vertical inhibition of the ERK-MAPK pathway, via concurrent MEK and RAF inhibition, is an approved therapeutic strategy for the treatment of melanoma (63), and concurrent inhibition of EGFR and BRAF is approved for the treatment of BRAF V600E mutant colorectal cancer (64). Therefore, we hypothesized that vertical inhibition of the autophagy pathway would result in enhanced growth suppression in autophagy-dependent PDAC.

### Concurrent inhibition of autophagosome nucleation and degradation enhances growth suppression

To elucidate whether pharmacological inhibition of VPS34 would phenocopy the results we observed via genetic depletion, we utilized the VPS34 inhibitor SAR405 (24) (henceforth referred to as VPS34i). First, we identified a dose range of VPS34i that led to an increase in the autophagy markers p62/*SQSTM1* and LC3B-II in our panel of PDAC cell lines (**Fig. 2A**; Supplementary Fig. S2A). As has been demonstrated in other cancer model systems (24,65), VPS34i treatment resulted in accumulation of both p62 and LC3B-II by 24 hours, and this increase was sustained at 72 hours of treatment (**Fig. 2A**; Supplementary Fig. S2A). To demonstrate that the accumulation of autophagy-related proteins was indicative of decreased flux through the autophagy pathway, we utilized a panel of PDAC cell lines that stably expressed mCherry-EGFP-LC3B, a fluorescent reporter that measures autophagic flux (58). We found that treatment with VPS34i (3 µM) for 24 hours resulted in a significant decrease in autophagic flux in the majority of cell lines tested (**Fig. 2B** and **C**; Supplementary Fig. S2B).

**Figure 2.**
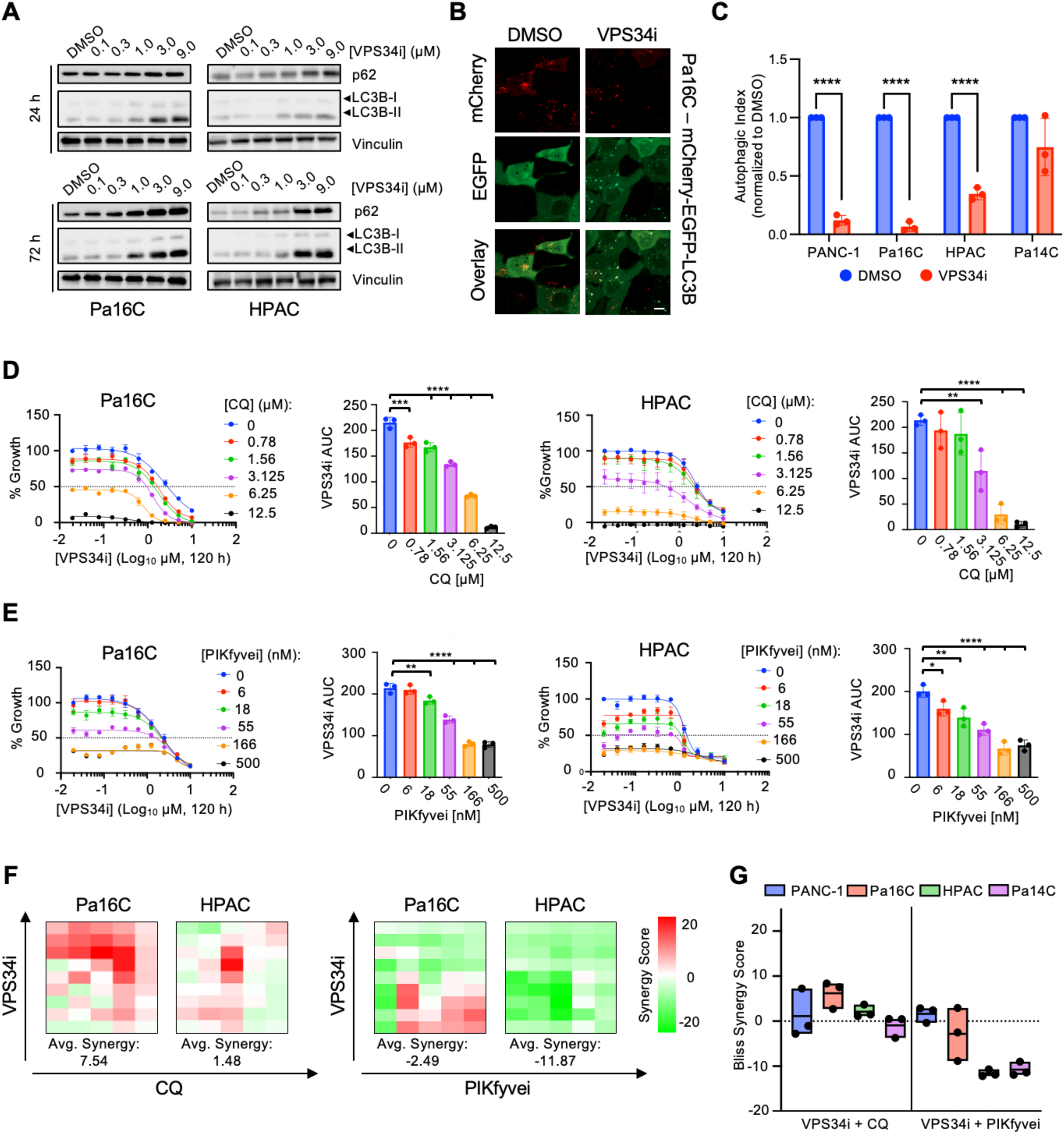
VPS34 inhibition sensitizes PDAC cells to inhibitors of the terminal stage of the autophagy pathway. **A,** Immunoblots of Pa16C and HPAC cell lines treated for 24 or 72 hours with VPS34i (SAR405) at indicated concentrations (representative of three independent experiments). **B,** Representative images of Pa16C mCherry-EGFP-LC3B cells treated with vehicle (DMSO) or VPS34i (SAR405, 3 µM) for 24 hours and imaged via confocal microscopy at 40X magnification. Scale bar, 20 µm. **C,** PDAC cells stably expressing mCherry-EGFP-LC3B were treated with vehicle or VPS34i (SAR405, 3 µM) and imaged by confocal microscopy. Data points are representative of the mCherry/EGFP ratio for each field (at least 9 fields per condition). Plot is representative of three independent experiments per cell line. Error bars represent mean ± SD of three independent experiments and statistical significance was determined by unpaired Student’s *t*-test. ****, π < 0.0001. **D,** Five-day viability assay in Pa16C and HPAC cells following increasing doses of VPS34i (SAR405, [0.039 – 10 µM]) alone or in combination with doses of CQ (chloroquine, [0.78 – 12.5 µM]) treatment. Each data point represents mean ± SEM of three independent experiments. Area under the curve (AUC) values from each biological replicate of data in proliferation curves are plotted, error bars denote mean ± SD. Statistical significance was determined by a one-way ANOVA with Dunnett’s test. **, π < 0.01; ***, π < 0.001; ****, π < 0.0001. **E,** Five-day viability assay in Pa16C and HPAC cells following increasing doses of VPS34i (SAR405, [0.039 – 10 µM]) alone or in combination with doses of PIKfyvei (apilimod, [6 – 500 nM]) treatment. Each data point represents mean ± SEM of three independent experiments. Area under the curve (AUC) values from each biological replicate of data in proliferation curves are plotted, error bars denote mean ± SD. Statistical significance was determined by a one-way ANOVA with Dunnett’s test. *, π < 0.05; **, π < 0.01; ****, π < 0.0001. **F,** Excess over bliss synergy scores calculated from biological replicate of data in **D** and **E** using SynergyFinder. Tiles represent synergy score for individual combination. **G,** Average synergy scores taken from studies in **D** and **E** and Supplementary Fig**. S2C, S4A**.

Given that genetic suppression of *PIK3C3* sensitized PDAC cells to CQ treatment, we hypothesized that combined VPS34i and CQ would more potently suppress cell growth. Indeed, we found that concurrent treatment with increasing doses of VPS34i and CQ reduced cellular proliferation across a panel of PDAC cell lines (**Fig. 2D**; Supplementary Fig. S2C). Extending our analysis to a 14-day clonogenic growth assay, we also observed greater growth suppression when combining VPS34i and CQ than when using either inhibitor alone (Supplementary Fig. S3A and B). We and others recently demonstrated that the lipid kinase phosphatidylinositol-3-phosphate 5-kinase (PIKfyve), which is integral to lysosomal function, is a targetable vulnerability in PDAC (29,30). Like CQ, the PIKfyve inhibitor apilimod impairs the terminal stage of the autophagy pathway (66). However, unlike CQ, apilimod (henceforth referred to as PIKfyvei) is a potent and specific pharmacological inhibitor (29). We found that concurrent treatment with increasing doses of VPS34i and PIKfyvei also reduced PDAC cellular proliferation in our panel of PDAC cell lines (**Fig. 2E**; Supplementary Fig. S4A). In the majority of cell lines tested, VPS34i synergistically enhanced CQ-mediated growth suppression, indicated by values >0 using Bliss synergy analysis (**Fig. 2F** and **G**; Supplemental Fig. S4B). However, combined VPS34i and PIKfyvei was not synergistic (**Fig. 2F** and **G**; Supplemental Fig. S4B). We hypothesize that this could be because PIKfyve is responsible for generating PI(3,5)P_2_ from PI3P (67,68), and PI3P is generated by VPS34 (23). This may limit the efficacy of combined PIKfyve and VPS34 inhibition.

### Vertical inhibition of autophagosome initiation and degradation enhances growth suppression

Having demonstrated that vertical inhibition of the nucleation and degradation stages of the autophagy pathway enhanced growth suppression in PDAC cell lines, we next asked whether this concept could be extended to the initiation phase. The ULK1/2 kinases are the human homologs of yeast *Atg1* and are crucial for autophagy initiation (69). In contrast to VPS34, which plays well-established roles in other endocytic processes such as endocytosis in addition to regulating autophagy (56,70), ULK1/2 are autophagy-specific kinases.

First, we validated that the ULK1/2 inhibitor ULK101 (26) (hereby referred to as ULKi) inhibited ULK1/2 function in our panel of human PDAC cell lines. To do this we immunoblotted for the activation of the ULK1/2 substrate ATG14 and found that phosphorylation of ATG14 decreased with increasing doses of ULKi, both at 4 and 24 hours (**Fig. 3A**; Supplementary Fig. S5A). Next, to determine whether ULKi treatment inhibited autophagosome formation, we expressed the fluorescent omegasome marker GFP-DFCP1 (71) in a panel of PDAC cell lines and monitored autophagosome formation via live cell microscopy. After treating with ULKi (3 µM) for 4 hours, we found that autophagosome formation was decreased across a panel of PDAC lines as measured by a decrease in GFP-positive puncta area per cell (**Fig. 3B and C**; Supplementary Fig. S5B).

**Figure 3.**
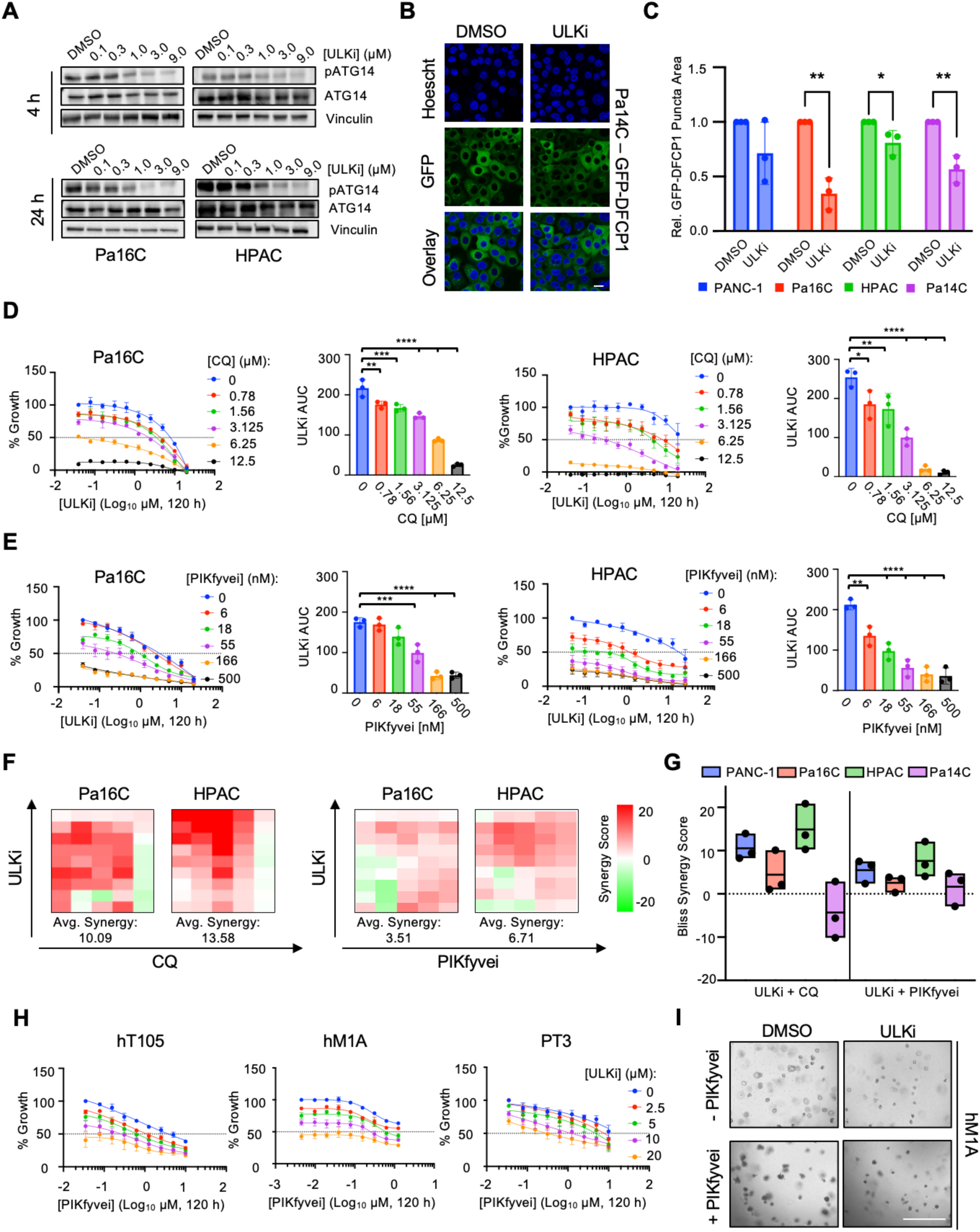
ULK1/2 inhibition impairs autophagosome formation and synergizes with CQ or PIKfyvei to decrease PDAC cell growth. **A,** Immunoblots of Pa16C and HPAC cell lines treated for 4 or 24 hours with ULKi (ULK101) at indicated concentrations (representative of three independent experiments). **B,** Representative images of Pa14C-GFP-DFCP1 cells treated with vehicle (DMSO) or ULKi (ULK101, 3 µM) for four hours and imaged via confocal microscopy at 40X magnification. Scale bar, 20 µm. **C,** PDAC cells stably expressing GFP-DFCP1 were treated with vehicle or ULKi (ULK101, 3 µM) and imaged by confocal microscopy. Data points are representative of the GFP area per cell ratio for each field (at least 9 fields per condition), normalized to DMSO control. Plot is representative of three independent experiments per cell line. Error bars represent mean ± SD of three independent experiments and statistical significance was determined by unpaired Student’s *t-*test. *, π < 0.05; **, π < 0.01. **D,** Five-day viability assay in Pa16C and HPAC cells following increasing doses of ULKi (ULK101, [0.078 – 20 µM]) alone or in combination with doses of CQ (chloroquine, [0.78 – 12.5 µM]) treatment. Each data point represents mean ± SEM of three independent experiments. Area under the curve (AUC) values are plotted from each biological replicate of data shown in proliferation curve, error bars denote mean ± SD. Statistical significance was determined by a one-way ANOVA with Dunnett’s test. *, π < 0.05; **, π < 0.01; ***, π < 0.001; ****, π < 0.0001. **E,** Five-day viability assay in Pa16C and HPAC cells following increasing doses of ULKi (ULK101, [0.078 – 20 µM]) alone or in combination with doses of PIKfyvei (apilimod, [6 – 500 nM]) treatment. Each data point represents mean ± SEM of three independent experiments. Area under the curve (AUC) values from each biological replicate of data plotted in proliferation curves are plotted, error bars denote mean ± SD. Statistical significance was determined by a one-way ANOVA with Dunnett’s test. **, π < 0.01; ***, π < 0.001; ****, π < 0.0001. **F,** Excess over bliss synergy scores calculated from biological replicate of data in **D** and **E** using SynergyFinder. Tiles represent synergy score for individual combination. **G,** Average synergy scores taken from studies in **D** and **E** and Supplementary Fig. **S5C** and **S7A**. **H,** Percent growth inhibition based on CellTiter-Glo 3D assay of human PDAC organoids treated with ULKi (ULK101) and PIKfyvei (apilimod) at indicated doses for 5 days. Data presented is the average of three biological replicates, error bars denote mean ± SD. **I**, Representative images of hM1A human organoids following treatment with DMSO, ULKi (ULK101, 20µM), PIKfyvei (apilimod, 1.25 µM) and ULKi + PIKfyvei for 5 days. Scale bar, 150 µm.

Next, we tested whether ULK1/2 inhibition sensitized PDAC cells to CQ-mediated growth suppression. Indeed, the combination of ULKi and CQ enhanced growth suppression in a 5-day proliferation assay (**Fig. 3D**; Supplementary Fig. S5C). Extending our analysis to a 14-day clonogenic growth assay, we also observed greater growth suppression when combining ULKi and CQ than when using either alone (Supplementary Fig. S6A and B). We found that concurrent treatment with increasing doses of ULKi and PIKfyvei similarly reduced PDAC cellular proliferation (**Fig. 3E**; Supplementary Fig. S7A). In the majority of cell lines tested, both combination treatments of ULKi and CQ or ULKi and PIKfyvei were synergistic (**Fig. 3F** and **G**; Supplementary Fig. S7B). Notably, ULKi-anchored combinations were more synergistic across cell lines than VPS34i-based combinations. Collectively, we found that multiple vertical autophagy inhibitor combinations reduced cell growth, with ULKi-based combinations demonstrating more synergy than VPS34i-based combinations.

Subsequently, we sought to determine whether vertical inhibition of the autophagy pathway would impact the growth of *KRAS*-mutant patient-derived organoid (PDO) models of PDAC. It is well appreciated that PDOs are more heterogeneous than 2D cell line models and that therapeutic profiles in organoids correspond more closely with patient outcomes (72,73). We treated PDOs with a dose response of ULKi and PIKfyvei alone and in combination and found that the combination further reduced cell viability (**Fig. 3H** and **I**; Supplementary Fig. 7C).

### Pathway profiling identifies signaling changes associated with vertical inhibition of the autophagy pathway

To better elucidate the mechanistic consequences of vertical inhibition of the autophagy pathway, we performed system-wide profiling of temporal signaling changes caused by single and multi-node autophagy inhibition using reverse-phase protein array (RPPA) analysis. We performed RPPA in a panel of four *KRAS*-mutant PDAC cell lines (PANC-1, MIA PaCa-2, Pa14C, and Pa02C) treated with vehicle, ULKi, CQ, PIKfyvei, ULKi + CQ, or ULKi + PIKfyvei for 4, 24, or 72 hours. We measured total and phosphoprotein levels of 139 different proteins spanning multiple cancer-related signaling pathways (Supplementary Table 3 and 4; Supplementary Fig. S8). Of note, we found that vertical inhibitor combinations (i.e. ULKi + CQ or ULKi + PIKfyvei) resulted in more significant signaling changes compared to vehicle than any single node inhibitor treatment (**Fig. 4A**). This indicates that vertical autophagy inhibition has more profound effects on cellular signaling than single node inhibition. Broadly, the signaling changes associated with vertical inhibition of the autophagy pathway were related to autophagy regulation, cell cycle progression, apoptosis, and the PI3K-mTOR pathway.

**Figure 4.**
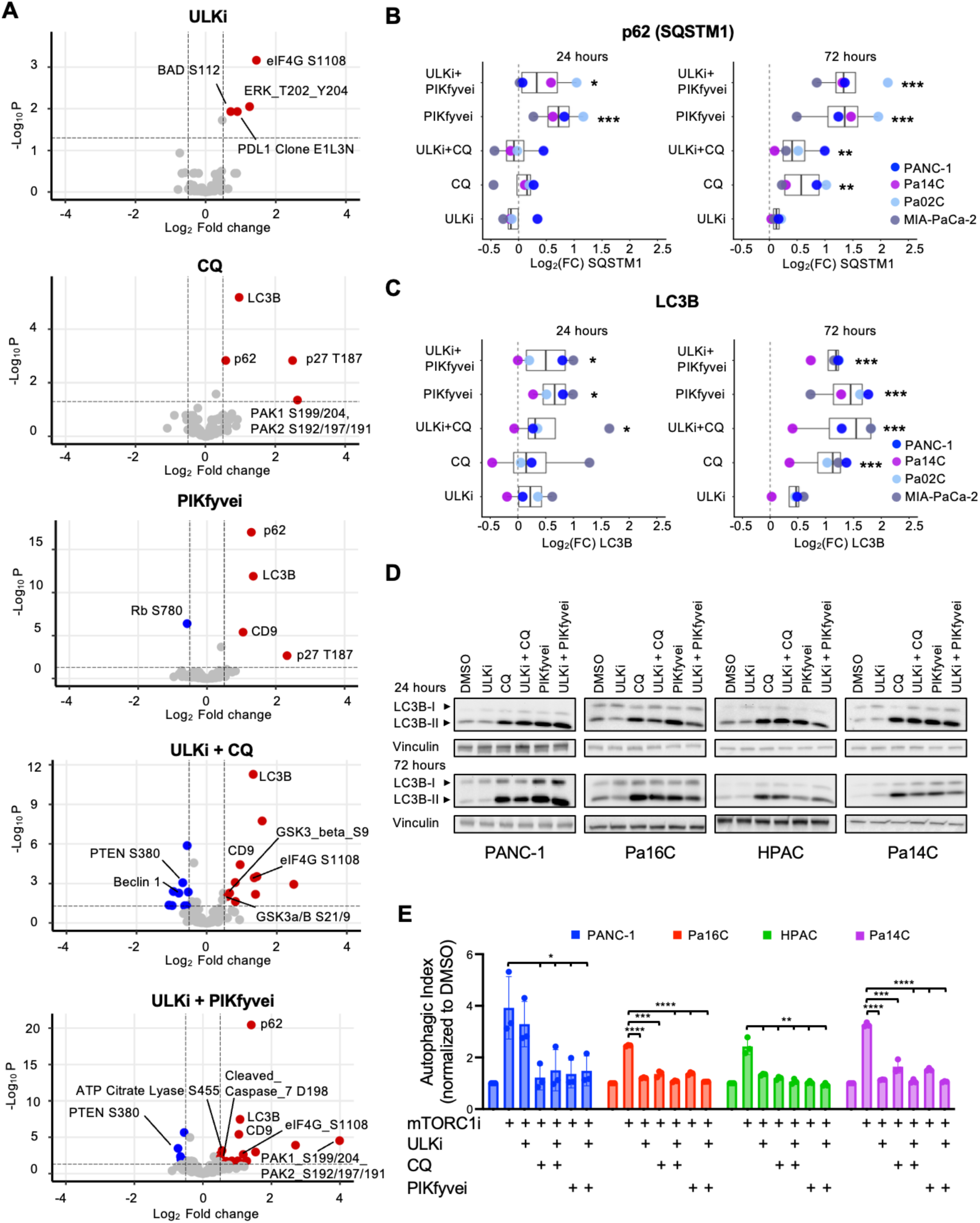
RPPA identifies signaling changes following single-agent and vertical autophagy inhibition. **A,** Volcano plots showing log2 fold-change (log2FC) on the X-axis and -Log_10_ adjusted p-values (-Log_10_ P) on the Y-axis. From RPPA experiment where PANC-1, Pa14C, Pa02C and MIA-PaCa-2 cells were treated with vehicle (DMSO), ULKi (ULK101, 3 µM), CQ (6.125 µM), PIKfyvei (apilimod, 100 nM), ULKi (ULK101) + CQ, or ULKi (ULK101) + PIKfyvei (apilimod) for 72 hours. Differential expression and significance were determined by linear modeling of all cell lines together and empirical Bayes moderation in limma. Shown are significantly (adj. p-val. < 0.05, dotted line) upregulated (in red) or downregulated (in blue) proteins across all cell lines. **B,** Log2FC values of p62/SQSTM1 antibody from RPPA experiment described in **A** compared to vehicle at 24 hours and 72 hours of treatment. Significance values calculated across all cell lines. *, π < 0.05; **, π < 0.01; ***, π < 0.001. **C,** Log2FC values of LC3B antibody from RPPA experiment described in **A** compared to vehicle at 24 hours and 72 hours of treatment. Significance values calculated across all cell lines. *, π < 0.05; ***, π < 0.001. **D,** Immunoblots of PANC-1, Pa16C, HPAC and Pa14C cell lines treated for 24 hours with vehicle (DMSO), ULKi (ULK101, 3 µM), CQ (6.125 µM), PIKfyvei (apilimod, 100 nM), ULKi (ULK101) +CQ (3 µM and 6.125 µM respectively), or ULKi (ULK101) + PIKfyvei (apilimod) (3 µM and 100 nM respectively) (representative of three independent experiments). **E,** Flow cytometry analysis of GFP-LC3-RFP-LC3ΔG-expressing PDAC cell lines following treatment with vehicle, mTORC1i (RMC-5552, 1 nM), ULKi (ULK101, 3 µM), CQ (6.25 µM), or PIKfyvei (apilimod, 100 nM) alone or in combination for 24 hours as denoted. The median values for GFP and RFP fluorescence for each condition were used to determine a ratio of RFP/GFP as a measure of autophagic flux which was then normalized to DMSO for each biological replicate. Data is representative of the median ± SD of three independent experiments. Statistical significance was determined by unpaired Student’s *t-*test. *, π < 0.05; **, π < 0.01; ***, π < 0.001; ****, π < 0.0001.

Consistent with inhibition of the autophagy pathway, the autophagy-related proteins LC3B/*MAP1LC3B* and p62/*SQSTM1* accumulated following both 24 and 72 hours of single agent CQ or PIKfyvei treatment (**Fig. 4B** and **C**). In contrast, single agent ULKi treatment did not result in an accumulation of LC3B or p62. However, the accumulation of LC3B and p62 was maintained when the autophagy pathway was inhibited using vertical inhibitor combinations (**Fig. 4B** and **C**).

LC3B-I is a ubiquitin-like protein that is modified to the mature LC3B-II form via the conjugation of a phosphatidylethanolamine (PE) by ATG proteins. This mature LC3B-II then integrates into forming autophagosomes and the accumulation of LC3B-II can be indicative of impaired autophagy (33). Given that the changes in the RPPA represent total LC3B and cannot be assigned to the immature LC3B-I or mature LC3B-II, we next evaluated LC3B accumulation via immunoblotting to assess the accumulation of LC3B-II and LC3B-I independently.

In alignment with our previous studies (29), we found that treatment with CQ or PIKfyvei led to an accumulation of LC3B-II at 24 hours which was sustained out to 72 hours of treatment indicative of autophagy inhibition (**Fig. 4D**). Conversely, when we treated with ULKi alone, we observed LC3B-I accumulation relative to DMSO at both 24 and 72 hours and did not observe consistent changes in LC3B-II protein levels (**Fig. 4D**). LC3B-I accumulation has been associated with ULK inhibition in other experimental systems (74), and this observation supports our previous conclusion that ULKi inhibits autophagy by decreasing autophagosome formation rather than by causing an accumulation of mature autophagosomes. Finally, in agreement with our findings in the RPPA study (**Fig. 4C**), we found that vertical autophagy inhibition also consistently caused increased levels of LC3B-II (**Fig. 4D**).

To validate and extend the changes in autophagic markers that we observed via RPPA-profiling and immunoblotting, we next assessed autophagic flux. We previously demonstrated that both CQ treatment and PIKfyve inhibition decreased autophagic flux in PDAC cell lines (29), and we initially hypothesized that vertical inhibition of the autophagy pathway would decrease flux more than single node inhibition. To test this hypothesis, we evaluated whether vertical inhibitor combinations could suppress autophagy induction more thoroughly than single node inhibition. To perform these assays, we treated with an inhibitor of mTORC1 (RMC-5552 (39), henceforth referred to as mTORC1i), as inhibition of mTORC1 is a standard method for inducing autophagy (33). We generated a panel of PDAC cell lines expressing the GFP-LC3-RFP-LC3ΔG autophagic flux probe (75) and utilized flow cytometry to assess autophagic flux. We treated cells with 1 nM of mTORC1i for 24 hours, a dose that inhibited phosphorylation of direct substrates of mTORC1 (S6 (S235/236) and 4EBP1 (T37/46)) at 24 hours (Supplementary Fig. S9 A). Indeed, we found that mTORC1i induced autophagic flux to 2-4-fold the basal level following 24 hours of treatment (**Fig. 4E**). We then treated cells with mTORC1i and autophagy inhibitors simultaneously and found that both single agent autophagy inhibitors and vertical combinations of autophagy inhibitors decreased mTORC1i-induced autophagic flux across our panel of PDAC cell lines (**Fig. 4E**). Notably, vertical autophagy inhibitor combinations did not decrease mTORC1i-induced autophagic flux to a greater degree than single node inhibition. We conclude that rather than further decreasing flux through the autophagic pathway when paired with CQ or PIKfyvei, inhibitors of upstream nodes of the autophagy pathway impact autophagosome formation (**Fig. 3B** and **C**). Therefore, autophagic flux is not further decreased by vertical autophagy inhibition compared to single node inhibition.

### Vertical inhibition of the autophagy pathway increases apoptosis

Our RPPA results also indicated that vertical inhibition of the autophagy pathway with concurrent ULKi and PIKfyvei led to the accumulation of several pro-apoptotic cleaved caspases (**Fig. 4A**; Supplementary Fig. S8). Of note, combined ULKi and PIKfyvei led to a significant accumulation of cleaved caspases 9 (D315), 7 (D198), and 6 (D162) while ULKi and CQ only led to a significant accumulation of cleaved caspase 9 (D315) (**Fig.5A**). We hypothesized that the accumulation of cleaved caspases may be indicative of increased apoptosis.

**Figure 5.**
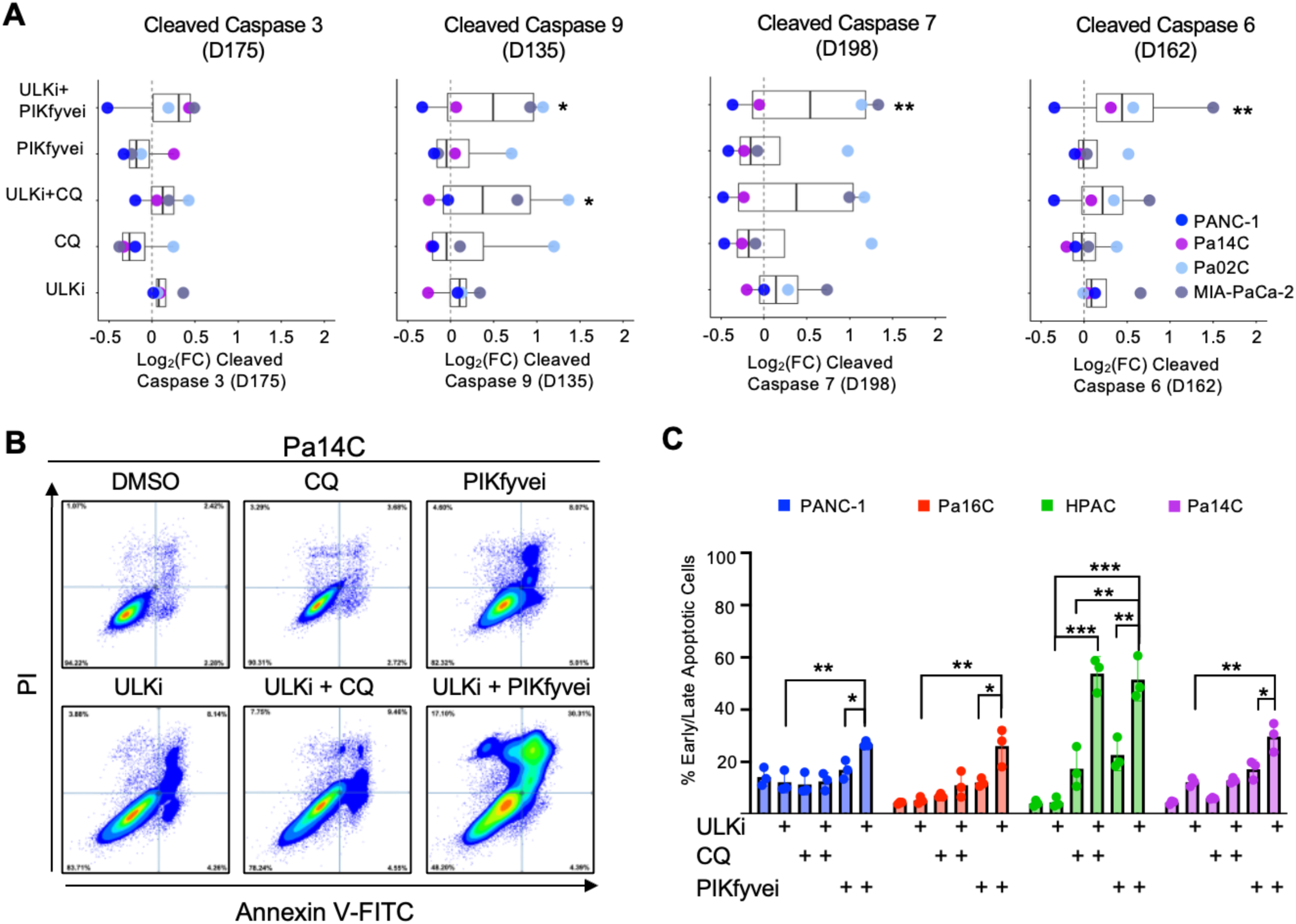
Vertical autophagy inhibition results in the accumulation of cleaved caspases and increased cell death. **A,** Log2FC values of listed antibodies from RPPA experiment described in Figure 4A compared to vehicle at 72 hours of treatment. Significance values calculated across all cell lines. *, π < 0.05; **, π < 0.01. **B,** Representative flow cytometry results following Annexin V-FITC and PI staining in Pa14C cells after 72 hours of treatment with vehicle (DMSO), ULKi (ULK101,3 µM), CQ (6.125 µM), PIKfyvei (apilimod, 100 nM), ULKi (ULK101) + CQ, or ULKi (ULK101) + PIKfyvei (apilimod). The sum of the percentage of cells in the lower right and upper right quadrants is the percentage of early and late apoptotic cells. **C,** Percentage of early and late apoptotic cells from 3 biological replicates of the experiment described in **B**. Error bars represent mean ± SD of three independent experiments and statistical significance was determined by unpaired Student’s *t-*test. *, π < 0.05; **, π < 0.01; ***, π < 0.001.

To elucidate whether these alterations in cellular signaling lead to increased apoptosis, we monitored Annexin V-FITC and propidium iodide (PI) staining with flow cytometry to quantify apoptotic cells. Indeed, we observed that vertical autophagy inhibition via ULKi and CQ induced an increase in early- and late-stage apoptotic cells when compared to either inhibitor alone in one PDAC cell line (HPAC), while vertical autophagy inhibition via ULKi and PIKfyvei induced greater apoptosis than either alone across the entire panel of PDAC lines tested (**Fig. 5B** and **C**). We speculate that the accumulation of cleaved caspases resulting from vertical autophagy inhibition ultimately leads to caspase-dependent apoptosis.

### Vertical inhibition of the autophagy pathway enhances dependency on PI3K-AKT-mTORC1 signaling

Our RPPA analysis showed significant differences in the activation of several members of the PI3K-AKT-mTORC1 signaling pathway following sustained vertical autophagy inhibition (72 hours) with either ULKi + PIKfyvei or ULKi + CQ (**Fig 4A**; Supplementary Fig. S8). We found that vertical autophagy inhibition via PIKfyvei and ULKi led to increased phosphorylation of AKT (T308), GSK-3α/β (S21, S9), 4E-BP1 (T37/T46) and EIF4G (S1108) (**Fig. 6A**; Supplementary Fig. S10A) which are all associated with increased signaling through the PI3K-AKT-mTORC1 signaling pathway (34,76). We also found that vertical autophagy inhibition led to decreased phosphorylation of PTEN at S380 which is a negative regulator of PI3K signaling (34), therefore serving as an additional marker of activation of this signaling cascade.

**Figure 6.**
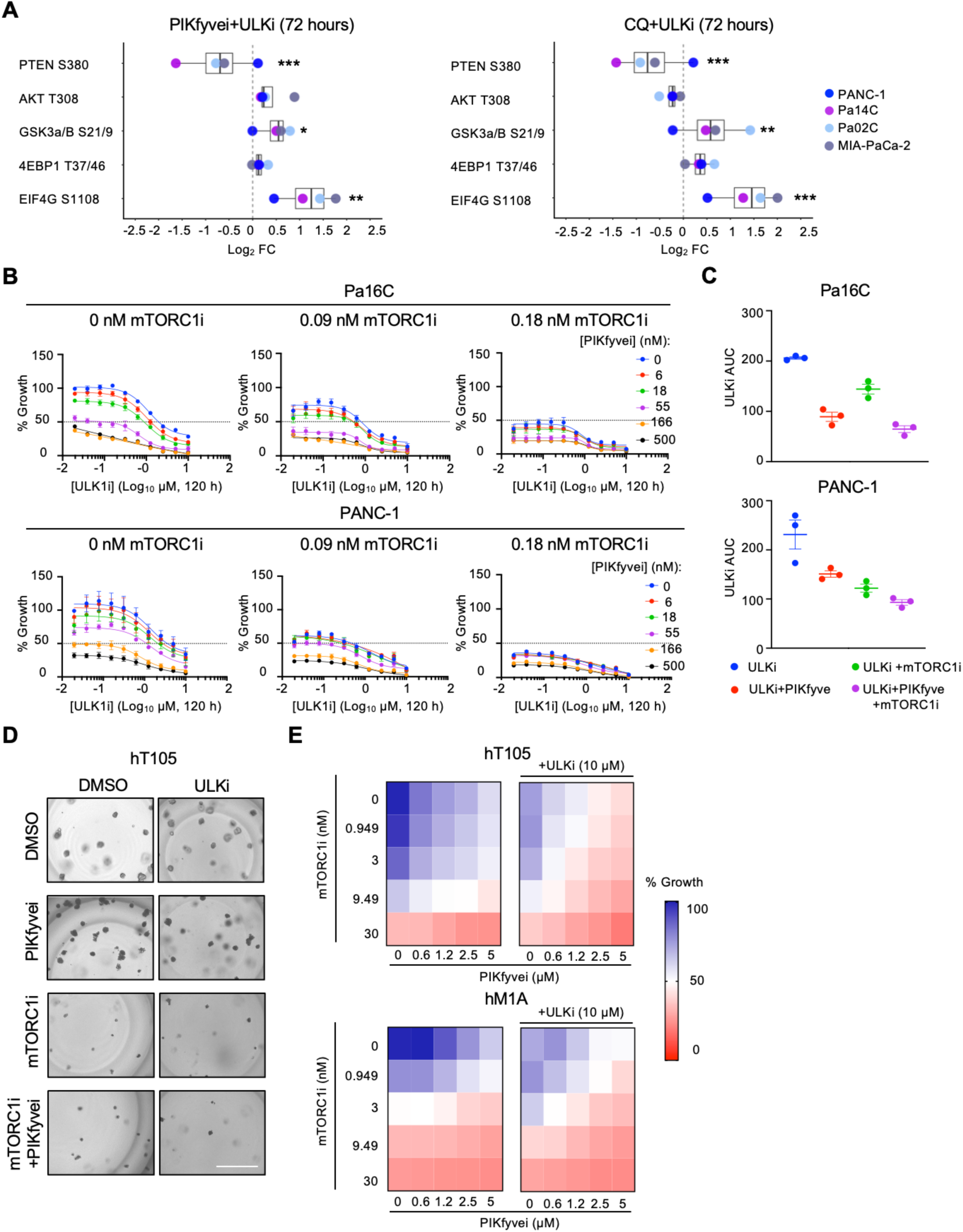
Vertical autophagy inhibition results in increased activation of PI3K-AKT-mTORC1 signaling pathway components resulting in enhanced sensitivity to mTORC1 inhibition. **A,** Log2FC values of listed antibodies from RPPA experiment described in Figure 4A compared to vehicle at 72 hours of treatment with ULKi + PIKfyvei (3 µM and 100 nM respectively) or ULKi + CQ (3 µM and 6.25 μM respectively). Significance values calculated across all cell lines. *, π < 0.05; **, π < 0.01; ***, π < 0.001. **B,** Five-day viability assay in Pa16C and PANC-1 cells following increasing doses of ULKi (ULK101, [0.039 – 10 µM]) alone or in combination with doses of PIKfyvei (apilimod, [6 – 500 nM]) treatment, with indicated dose of mTORC1i (RMC-5552). Each data point represents mean ± SEM of three independent experiments. **C,** Area under the curve (AUC) values from each biological replicate of ULKi data plotted in **B**, alone or in combination with 0.09 nM mTORC1i (RMC-5552) and 55 nM PIKfyvei (apilimod), error bars denote mean ± SD. **D,** Representative images of human hT105 organoids following treatment with mTORC1i (RMC-5552) and PIKfyvei (apilimod) at indicated doses +/- 10 µM of ULK101 for 5 days. Scale bar, 150 µm. **E,** Percent growth inhibition based on CellTiter-Glo 3D assay of human PDAC organoids treated with mTORC1i (RMC-5552) and PIKfyvei (apilimod) at indicated doses +/- 10 µM of ULK101 for 5 days. Data presented is the average of three biological replicates.

Taken together, these alterations in PI3K-AKT-mTORC1 signaling led us to hypothesize that inhibition of the autophagy pathway leads to altered metabolic sensing that increased PI3K-AKT-mTORC1 signaling. Consistent with this hypothesis, *MTOR* and *RPTOR*, both members of the mTORC1 complex, were significant hits in our CQ-anchored CRISPR-Cas9 mediated loss-of-function screen (**Fig. 1B**). Previous studies established that mTOR inhibition can trigger protective autophagy in cancer cells and mTOR inhibition can enhance sensitivity to autophagy inhibition (77). Recently, a bi-steric mTOR complex 1 (mTORC1)-selective inhibitor, RMC-5552 (mTORC1i) has been developed (39) and demonstrated tolerability and selective inhibition of mTORC1 in patients (NCT04774952) (41). We treated a panel of PDAC cells with mTORC1i and found that it potently inhibited cell growth (Supplemental Fig. S10B). To elucidate whether vertical autophagy inhibition sensitized pancreatic cancer cells to pharmacological inhibition of mTORC1, we performed 5-day cellular proliferation assays in PDAC cells following concurrent treatment with ULKi, PIKfyvei and mTORC1i (**Fig. 6B**). We found that the addition of mTORC1i potentiated the antiproliferative effects of concurrent ULKi and PIKfyvei as indicated by a decrease in the AUC in the triple combination compared to ULKi, ULKi + PIKfyvei or mTORC1i + ULKi (**Fig. 6C**). Furthermore, we also found that concurrent treatment with increasing doses of mTORC1i and ULKi reduced PDAC cellular proliferation (Supplementary Fig. S11A), and that in the majority of cell lines tested, treatment with mTORC1i and ULKi was synergistic (Supplementary Fig. S11B and C). The addition of PIKfyvei potentiated the antiproliferative effects of the mTORC1i + ULKi combination (Supplemental Fig. S11D). Next, we evaluated this combination in PDAC PDOs and found that the triple combination robustly suppressed growth (**Fig. 6D** and **E**). We conclude that vertical autophagy inhibition upregulates mTORC1 signaling as a resistance mechanism to loss of autophagy, leading to enhanced sensitivity to mTORC1 inhibition.

## DISCUSSION

Despite well characterized genetic drivers of PDAC, for most patients there are currently no approved targeted therapies (1–3). PDAC has a well-established dependency on autophagy for tumor growth (6,10,11). Unfortunately, clinical trials to test the efficacy of hydroxychloroquine (HCQ), the sole FDA-approved inhibitor of autophagy, as a single agent or in combination with other established PDAC therapies, have been largely unsuccessful in part due to HCQ lacking potency and efficacy (15–18). In this study, we identified and validated that concurrent inhibition of multiple nodes of the autophagy pathway more effectively reduced cellular proliferation. Our analysis of this combination strategy revealed that vertical inhibition of the autophagy pathway reduced autophagic flux while also leading to enhanced apoptotic cell death. Finally, we determined that vertical autophagy inhibition resulted in metabolic rewiring, leading to an increased dependence on PI3K-AKT-mTORC1 signaling that could be therapeutically exploited. This study elucidates how vertical inhibition of the autophagy pathway impacts cellular signaling and growth. Furthermore, our findings provide preclinical support for combining vertical autophagy inhibitor combinations with mTORC1 inhibition to further impede PDAC growth and enhance the efficacy of autophagy inhibition as a therapeutic strategy.

We demonstrated that genetic suppression of *PIK3C3*, the gene that encodes VPS34, sensitized PDAC cells to CQ treatment. VPS34 is a lipid kinase that converts PIP to PI3P, which is essential for the nucleation of the autophagosome (23,56). This lipid then recruits effectors important for the formation of the autophagosome, including double FYVE domain-containing protein 1 (DFCP1) and WIPI family members, thereby potentiating the generation of the autophagosome (56,78). Inhibition of VPS34 via the preclinical compound SAR405 has been shown to decrease growth of both colorectal and melanoma cancer models (24,65), but these findings have not been demonstrated in PDAC. In this study, we extend these findings to PDAC. We validated that pharmacological inhibition of VPS34 decreased autophagic flux in PDAC cells and reduced PDAC cell growth. Importantly, growth was further suppressed by the addition of CQ or PIKfyvei. While preclinical cancer studies have proposed combinations of VPS34 inhibitors with immunotherapies and mTOR inhibition (24,65), our studies are the first to propose combining a VPS34 inhibitor with an inhibitor of the degradation stage of the autophagy pathway. Thus far, VPS34 inhibition has not been pursued clinically. It is worth noting that VPS34 can form other complexes that are important for endocytosis and cellular trafficking (56,70). Given that VPS34 is also important for endocytic processes outside of the autophagy pathway, it is possible that this protein is essential to normal cellular functions and therefore may not be a suitable clinical target. Whereas VPS34 may not currently be a clinically tractable target, these findings led us to explore other, more translatable vertical inhibition strategies for targeting the autophagy pathway.

ULK1/2 are the initiating kinases of the autophagy pathway. While ULK1/2 inhibition has been previously characterized as a potential strategy for inhibiting autophagy in pancreatic cancer (22) and other cancer types, including non-small cell lung cancer (NSCLC) (28), this strategy has been historically limited by a lack of therapeutically relevant ULK inhibitors. Recently, an ULK1/2 inhibitor, DCC-3116, has entered clinical evaluation (NCT04892017, NCT05957367). Similar to what has been demonstrated using ULK inhibitors in other cell types (26), we validated that ULK1/2 inhibition decreased autophagosome formation. As this is a more therapeutically relevant and potentially autophagy-specific target than VPS34, we sought to determine if ULK1/2 inhibition would sensitize PDAC cells to CQ-mediated decreases in cell growth. Indeed, we found that ULK1/2 inhibition potentiated CQ- or PIKfyvei-mediated growth suppression.

We and others recently demonstrated that inhibition of PIKfyve, a lipid kinase responsible for converting PI3P to PI(3,5)P2, which is essential for autophagosome fission and fusion, is a potent method for inhibiting the degradation stage of the autophagy pathway and impairs PDAC growth in both *in-vitro* and *in-vivo* models (29,30). Based on these findings, we sought to elucidate whether inhibition of autophagy at the initiation stage (via an ULK1/2 inhibitor) or the nucleation stage (via a VPS34 inhibitor) would sensitize cells to PIKfyve inhibition in place of CQ treatment. We found that similar to concurrent ULK1/2 or VPS34 inhibition with CQ, ULK1/2 or VPS34 inhibition sensitized PDAC cells to PIKfyve inhibition mediated growth inhibition across a panel of PDAC cell lines. Therefore, we demonstrated that via multiple chemically distinct approaches, vertical inhibition of the autophagy pathway decreased PDAC cell growth.

We previously demonstrated that PIKfyvei is a more durable suppressor of PDAC growth than CQ treatment (29). This study provided further mechanistic insights regarding the effect of CQ and PIKfyvei on cell death signaling, particularly within the context of vertical inhibition of the autophagy pathway. We found that vertical autophagy inhibition via PIKfyvei and ULKi led to increased levels of multiple cleaved caspases. We also demonstrated that these increased levels of cleaved caspases were indicative of increased apoptosis. Vertical autophagy inhibition via ULKi and PIKfyvei induced greater apoptosis than either alone across the entire panel of PDAC lines tested, whereas the ULKi + CQ combination only induced more apoptosis than single agent treatment in one of the four cell lines. These observations support the ongoing development of improved PIKfyvei inhibitors (30,79), some of which are currently under clinical evaluation (NCT5988918).

Autophagy inhibition affects tumor extrinsic signaling within the tumor microenvironment, specifically immune cell signaling (65,80–84). Autophagy inhibition has been shown to enhance anti-tumor T-cell responses in pancreatic cancer, thereby enhancing sensitivity to immunotherapy (81). Selective inhibition of VPS34 by SAR405 has been demonstrated to enhance sensitivity to anti PD-1/PD-L1 immunotherapy (65). Additionally, inhibition of PIKfyve has been shown to enhance sensitivity to immune checkpoint blockade in prostate cancer models (80). We observed increased expression of immune markers following autophagy inhibition in our RPPA profiling. It will be important for future studies to determine the impact of vertical autophagy inhibition on the tumor microenvironment, especially as more autophagy inhibitors enter the clinic for pancreatic cancer treatment.

Autophagy inhibition in PDAC has been demonstrated to lead to alterations in other metabolic processes, including mitochondrial function (85) and dependence on fatty acid synthesis (30). In alignment with this, our RPPA based protein pathway activation mapping resulted in the identification of significant alterations in the activation/expression of several proteins related to metabolic signaling pathways following concurrent ULK and PIKfyve inhibition. This included activation of ATP Citrate lyase (S455), which is associated with increased fatty acid synthesis (86) and increased CD9 expression, which has been demonstrated to play a role in enhanced glutamine uptake (87). Additionally, we observed metabolic rewiring following vertical autophagy inhibition, resulting in enhanced signaling through the PI3K-AKT-mTORC1 pathway and increased activation of PAK following vertical autophagy inhibition, which we speculate could be indicative of increased usage and/or dependence on other nutrient scavenging pathways such as macropinocytosis (9,88–91) following prolonged inhibition of autophagy. These observations support more comprehensive metabolic studies in the future to elucidate the impact of vertical autophagy inhibition on global metabolism.

The interplay between the autophagy pathway and mTOR signaling is well appreciated as mTORC1 directly phosphorylates both ULK1 and VPS34 to suppress the initiation of autophagy (51,92). Our mechanistic studies revealed that PI3K-AKT-mTORC1 signaling was enhanced following prolonged vertical inhibition of the autophagy pathway, indicative of metabolic rewiring. This expands upon previous studies, which have demonstrated that inhibition of PIKfyve in PDAC leads to increased activation of mTOR signaling (30). We extended these studies by demonstrating that growth suppression by vertical inhibition of the autophagy pathway in PDAC was further enhanced by the addition of an mTORC1 inhibitor in both cell culture and organoid models. While others have shown that combined inhibition of mTORC1 and CQ treatment decreased proliferation and enhanced cell death in PDAC (93), this is the first study to demonstrate that vertical inhibition of the autophagy pathway in combination with mTORC1 inhibition decreases PDAC growth in both 2D and 3D models. mTORC1 is a promising potential therapeutic target in PDAC, and the mTORC1-specific inhibitor utilized in this study (RMC-5552) showed an encouraging safety and efficacy profile in a Phase I trial that included a pancreatic cancer patient (41). These observations support the importance of future *in vivo* studies to further elucidate the impacts of the combination strategy in *in vivo* models of PDAC. In summary, our studies identified and validated a novel method of inhibiting the autophagy pathway, via vertical inhibition, in autophagy-driven PDAC and provided a detailed molecular characterization of signaling changes following vertical autophagy inhibition. Our findings support combined vertical autophagy inhibition and mTORC1 inhibition as a future potential therapeutic strategy for the treatment of PDAC.

## ACKNOWLEDGEMENTS

We thank A. Maitra (MD Anderson Cancer Center) for PDAC cell lines, and D. Tuveson (Cold Spring Harbor Laboratory) and C. Kuo (Stanford) for organoids. We thank K. Wood (Duke) for the CRISPR-Cas9 druggable genome library. K.L. Bryant was supported by grants from the Pancreatic Cancer Action Network/AACR (15-70-25-BRYA), the Pancreatic Cancer Action Network (22-WG-DERB), National Cancer Institute (P50CA257911, R37CA251877, and R01CA262414), the Department of Defense (W81XWH2110693), and the Hirshberg Foundation (HF-2025-113). E.F. Petricoin is a shareholder and consultant of Ignite Therapeutics, Inc. DAiNA, Inc., and Perthera, Inc. E.F. Petricoin has received funding support from Mirati Therapeutics, Springworks Therapeutics, Inc, Deciperha, Inc., Genentech, Inc., and Abbvie, Inc. C.A. Stalnecker was supported by grants from the National Cancer Institute (T32CA009156 and F32CA232529). M.K. Roach was supported by a grant from the National Institute of General Medical Sciences (T32GM135095). R. Robb was supported by a grant from the National Cancer Institute (T32CA071341 and F31CA284869). The Microscopy Services Laboratory was supported by a Cancer Center Core Support grant to the UNC Lineberger Comprehensive Cancer Center (P30 CA016086).

## AUTHOR CONTRIBUTIONS

**M.K. Roach:** Conceptualization, formal analysis, investigation, visualization, methodology, validation, writing-original draft, writing-review, and editing. **S.E. Degan:** Formal analysis, investigation, visualization, methodology, writing-review, and editing. **J.M. DeLiberty**: Conceptualization, formal analysis, investigation, methodology. **L.M. Pita:** Formal analysis, investigation, writing-original draft. **N.L. Pieper:** Formal analysis and investigation. **R. Yang:** Formal analysis and investigation. **K. E. Taylor:** Formal analysis and investigation. **E.G. Schechter:** Formal analysis and investigation. **R. Robb:** Formal analysis and investigation. **M. Pierobon:** Formal analysis and investigation. **C.A. Stalnecker:** Formal analysis and investigation. **E.F. Petricoin:** Resources, supervision, writing-review and editing, and funding acquisition. **K.L. Bryant:** Conceptualization, formal analysis, resources, supervision, funding acquisition, project administration, writing-original draft, writing-review, and editing.

