## Supplementary Figures and Legends for "Vertical inhibition of the autophagy pathway impairs growth and enhances sensitivity to mTORC1 inhibition in pancreatic ductal adenocarcinoma"

#### Supplementary Fig. S1

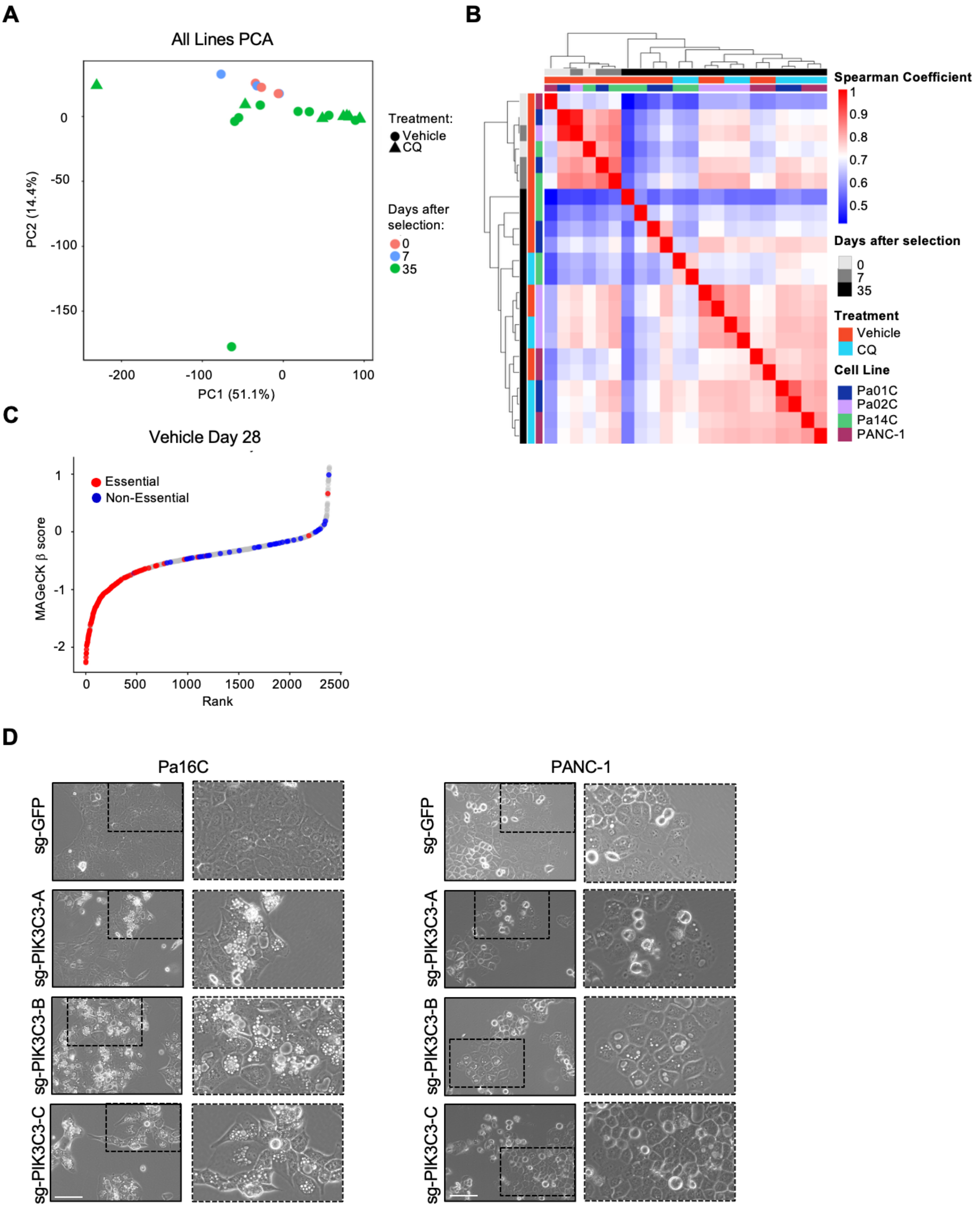

**Supplementary Figure 1**

CRISPR Cas9 loss-of-function screen identifies genetic modulators to CQ treatment. **A,** Principal component analysis (PCA) of median normalized sgRNA collected at 0, 7 and 35-days post-selection for all cell lines **B,** Spearman correlation for sgRNA collected at 0, 7, and 35-days post-selection for all cell lines in vehicle and CQ treatment conditions. **C,** Snake plot of the relative abundance of sgRNAs in the 28-day vehicle condition considered essential or non-essential by DepMap in pancreatic cancer cell lines. **D,** Representative wide field images of PANC-1 and Pa16C cells infected with control guide (sg-GFP) or three distinct sgRNAs against *PIK3C3* (sg-PIK3C3-A, sg-PIK3C3-B, sg-PIK3C3-C). Scale bar, 150 μm.

#### Supplementary Fig. S2

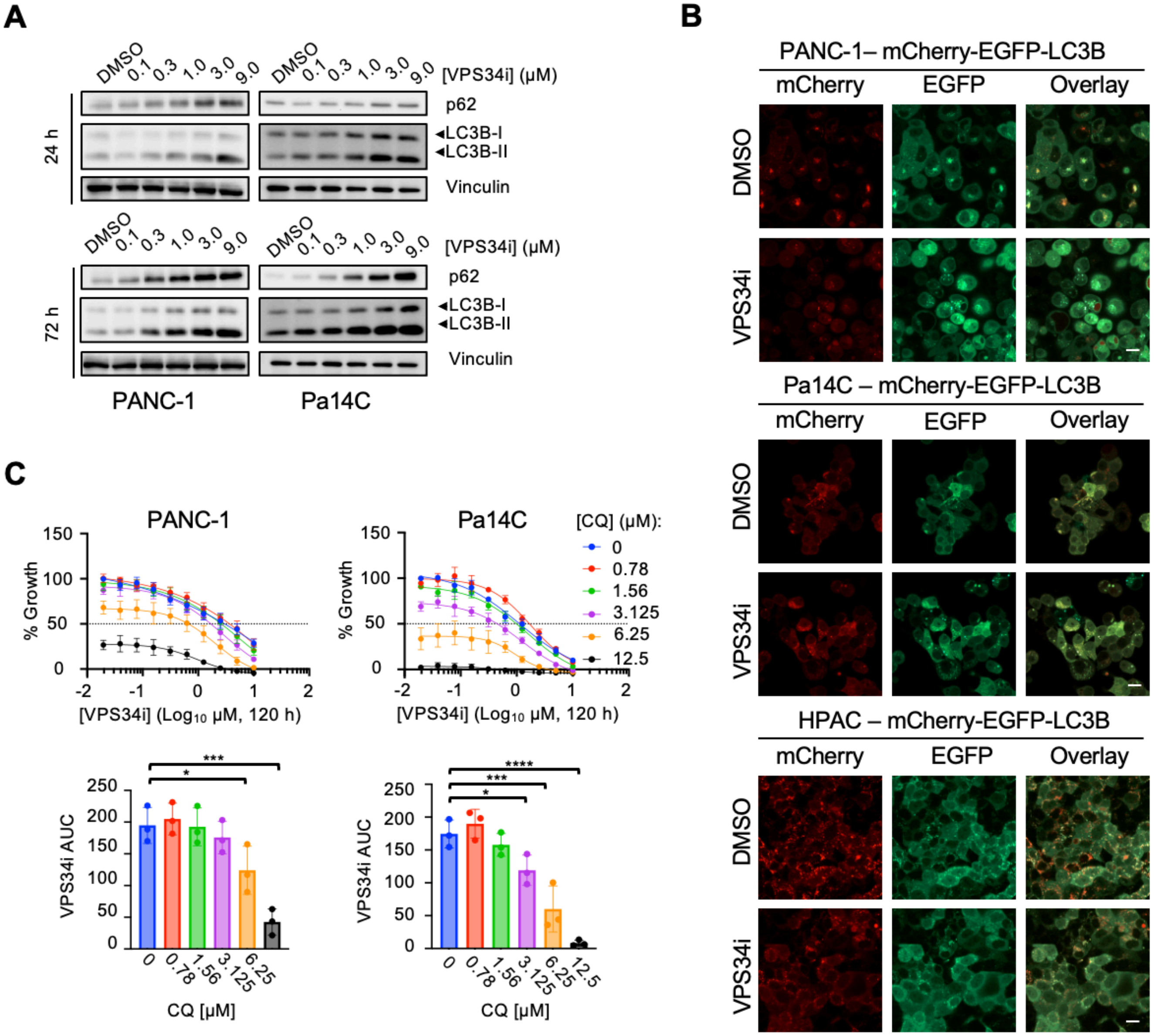

**Supplementary Figure 2**

PDAC cells treated with VPS34 inhibitor decrease autophagic flux and are sensitized to CQ treatment. **A,** Immunoblots of PANC-1 and Pa14C cell lines treated for 24 or 72 hours with VPS34i (SAR405) at indicated concentrations (representative of three independent experiments). **B,** Representative images for PANC-1, HPAC and Pa14C -mCherry-EGFP-LC3B cells treated with DMSO or VPS34i (SAR405, 3 μM) described and quantified in Fig. **2B-C**. Scale bar, 20μm. **C,** Top, five-day viability assay in PANC-1 and Pa14C cells following increasing doses of VPS34i (SAR405, [0.039 – 10 μM]) alone or in combination with doses of CQ (chloroquine, [0.78 – 12.5 μM]) treatment. Each data point represents mean ± SEM of three independent experiments. Bottom, area under the curve (AUC) values from each biological replicate of data plotted above, error bars denote mean ± SD. Statistical significance was determined by a one-way ANOVA with Dunnett's test. *, $\rho < 0.05$; ***, $\rho < 0.001$; ****, $\rho < 0.0001$.

#### Supplementary Fig. S3

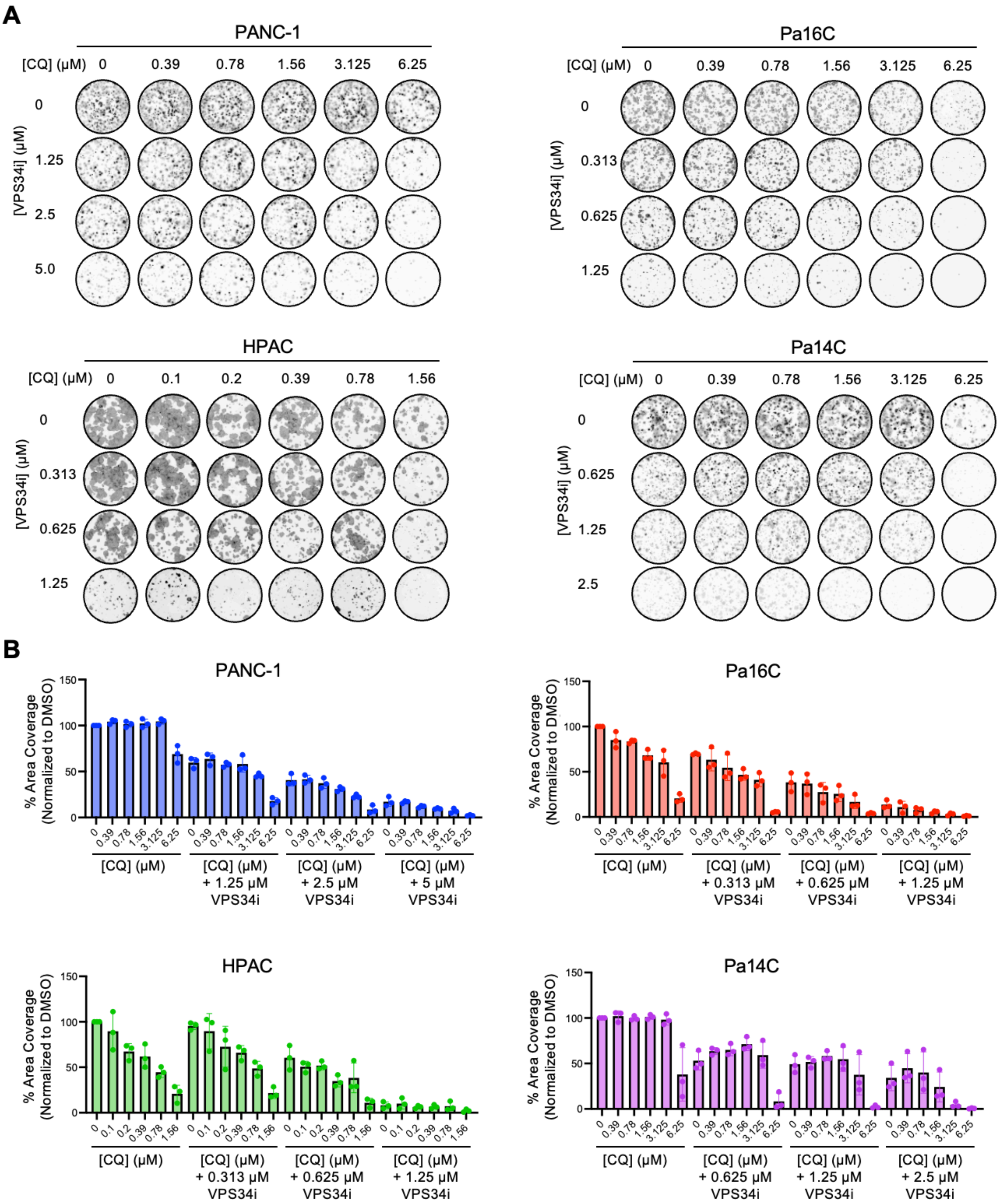

**Supplementary Figure 3**

VPS34 inhibition sensitizes cells PDAC cells to CQ following prolonged treatment. **A,** 10-14 day colony formation assay in PDAC cells treated with DMSO, or a range of doses of CQ, alone and in combination with a range of concentrations of VPS34i (SAR405) (representative of three independent experiments). **B,** Quantification of the average area coverage from the colony formation assays in **A**. Error bars denote mean ± SD of three independent experiments.

#### Supplementary Fig. S4

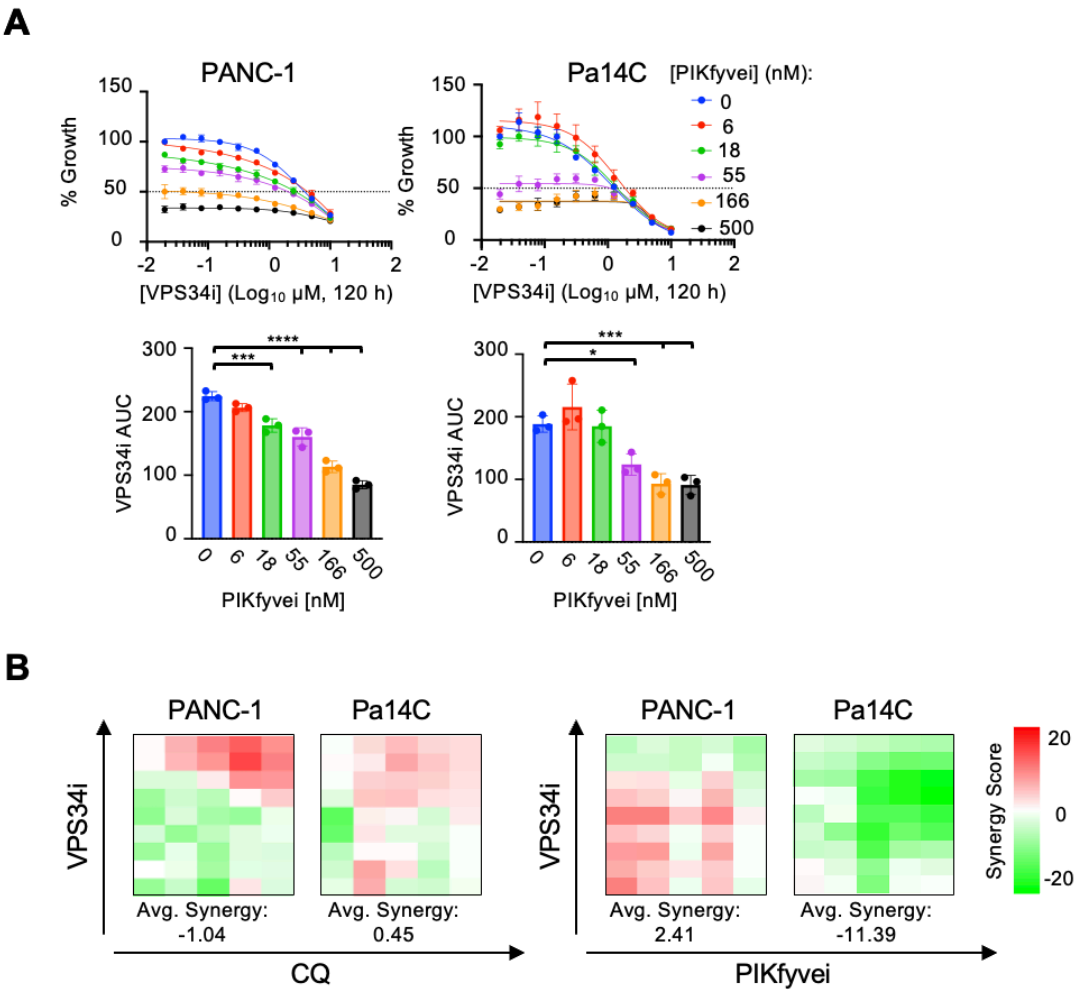

**Supplementary Figure 4**

VPS34 inhibition sensitizes cells PDAC cells to PIKfyvei. **A,** Top, five-day viability assay in PANC-1 and Pa14C cells following increasing doses of VPS34i (SAR405, [0.039 – 10 μM]) alone or in combination with doses of PIKfyvei (apilimod, [6 – 500 nM]) treatment. Each data point represents mean ± SEM of three independent experiments. Bottom, area under the curve (AUC) values from each biological replicate of data plotted above, error bars denote mean ± SD. Statistical significance was determined by a one-way ANOVA with Dunnett's test. *, $\rho < 0.05$; ***, $\rho < 0.001$; ****, $\rho < 0.0001$. **B,** Excess over bliss synergy scores calculated from biological replicate of data in **S2C** and **S4A** using SynergyFinder. Tiles represent synergy score for individual combination.

#### Supplementary Fig. S5

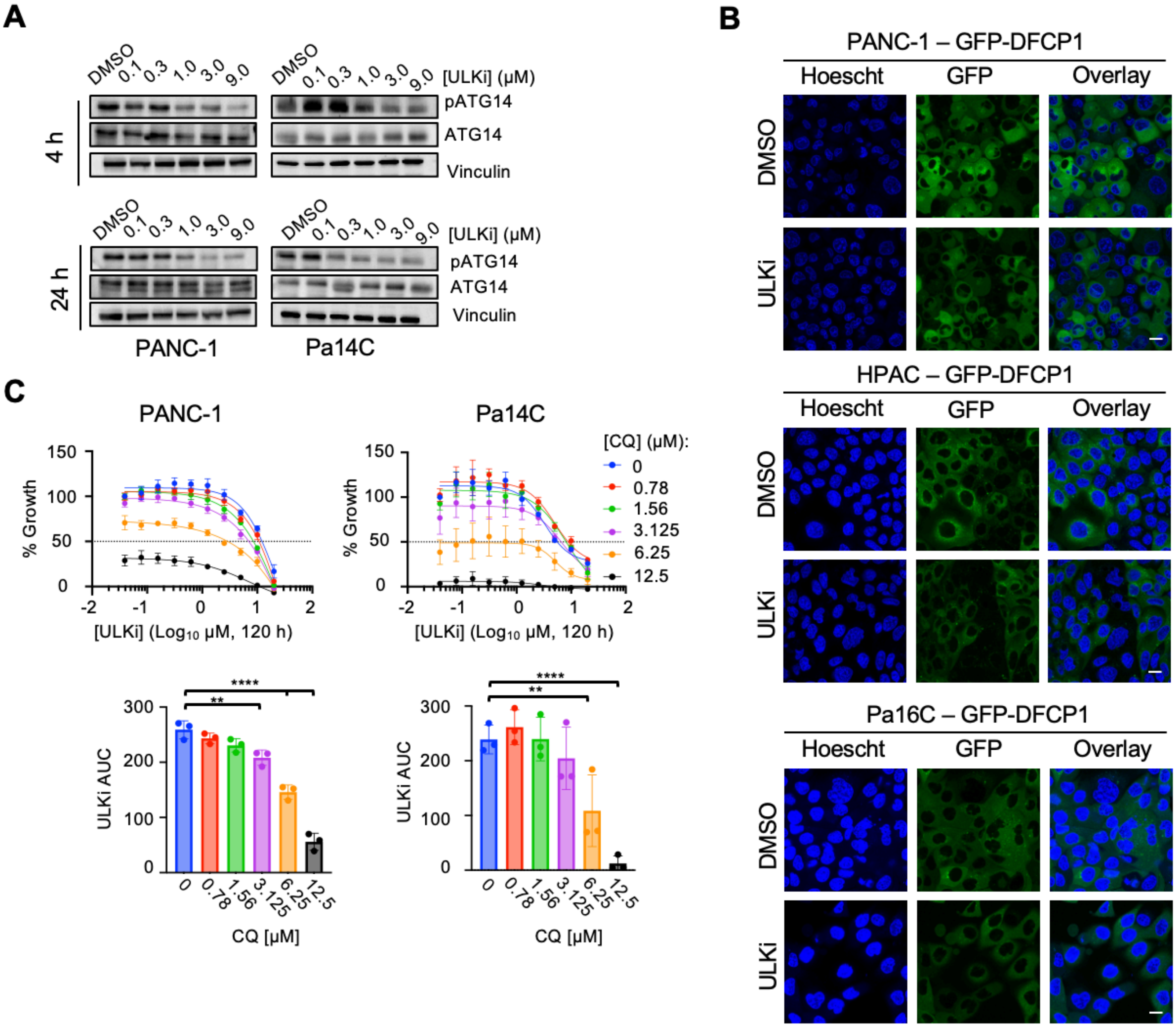

**Supplementary Figure 5**

ULKi decreases autophagosome formation and sensitizes PDAC cells to CQ treatment. **A,** Immunoblots of PANC-1 and Pa14C cell lines treated for 4 or 24 hours with ULKi (ULK101) at indicated concentrations (representative of three independent experiments). **B**, Representative images for PANC-1, HPAC, and Pa16C-GFP-DFCP1 cells treated with DMSO or ULKi (ULK101, 3 μM) described and quantified in Fig **3B-C**. Scale bar, 20μm. **C,** Top, five-day viability assay in PANC-1 and Pa14C cells following increasing doses of ULKi (ULK101, [0.078 – 20 μM]) alone or in combination with doses of CQ (chloroquine, [0.78 – 12.5 μM]) treatment. Each data point represents mean ± SEM of three independent experiments. Bottom, area under the curve (AUC) values from each biological replicate of data plotted above, error bars denote mean ± SD. Statistical significance was determined by a one-way ANOVA with Dunnett's test. **, $\rho < 0.01$; ****, $\rho < 0.0001$.

### Supplementary Fig. S6

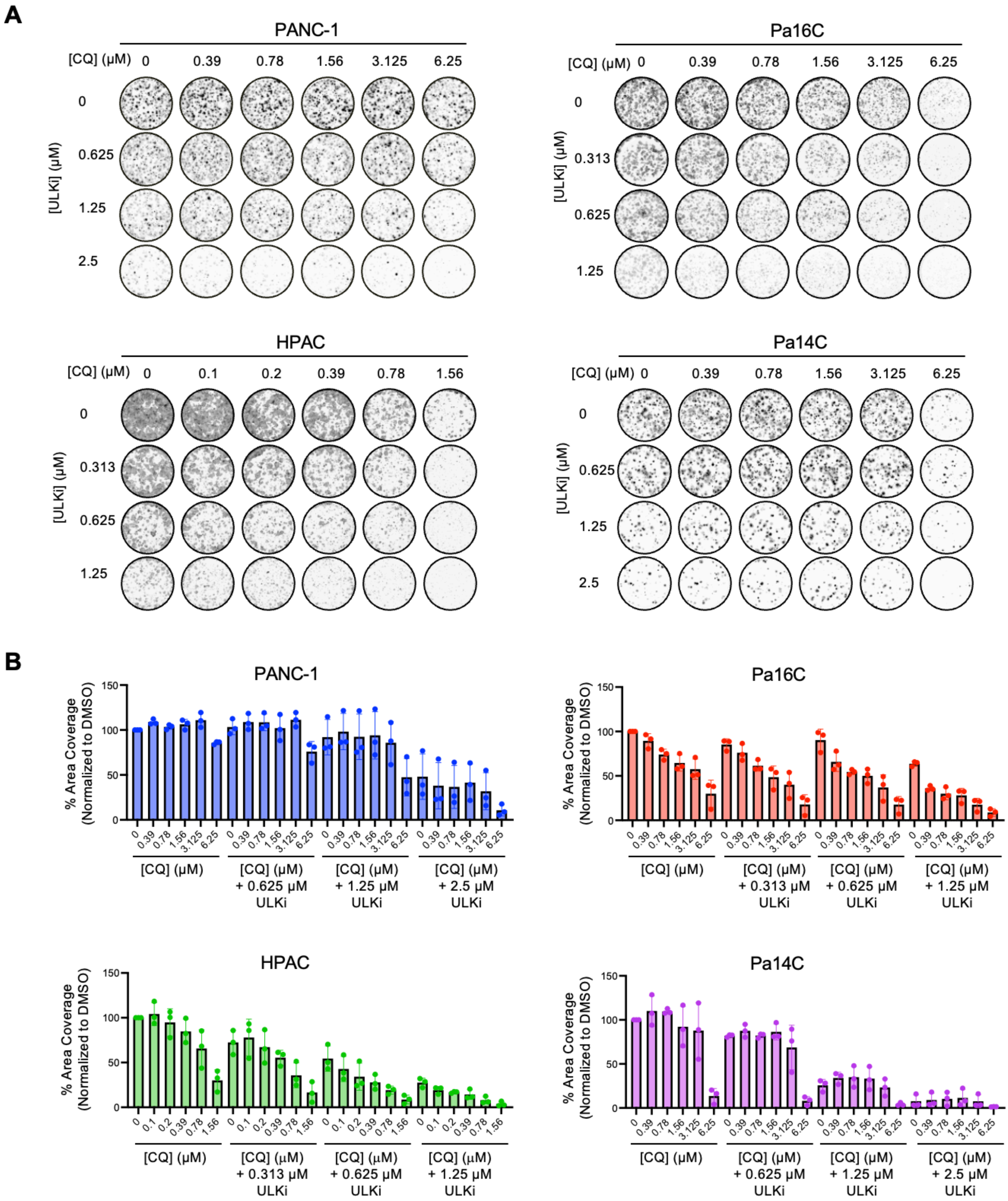

**Supplementary Figure 6**

ULK1/2 inhibition sensitizes PDAC cells to CQ treatment in colony formation assay. **A,** 10-14 day colony formation assay in PDAC cells treated with DMSO, or a range of doses of CQ alone and in combination with a range of concentrations of ULKi (ULK101) (representative of three independent experiments). **B,** Quantification of the average area coverage from the colony formation assays in **A**. Error bars denote mean ± SD of three independent experiments.

### Supplementary Fig. S7

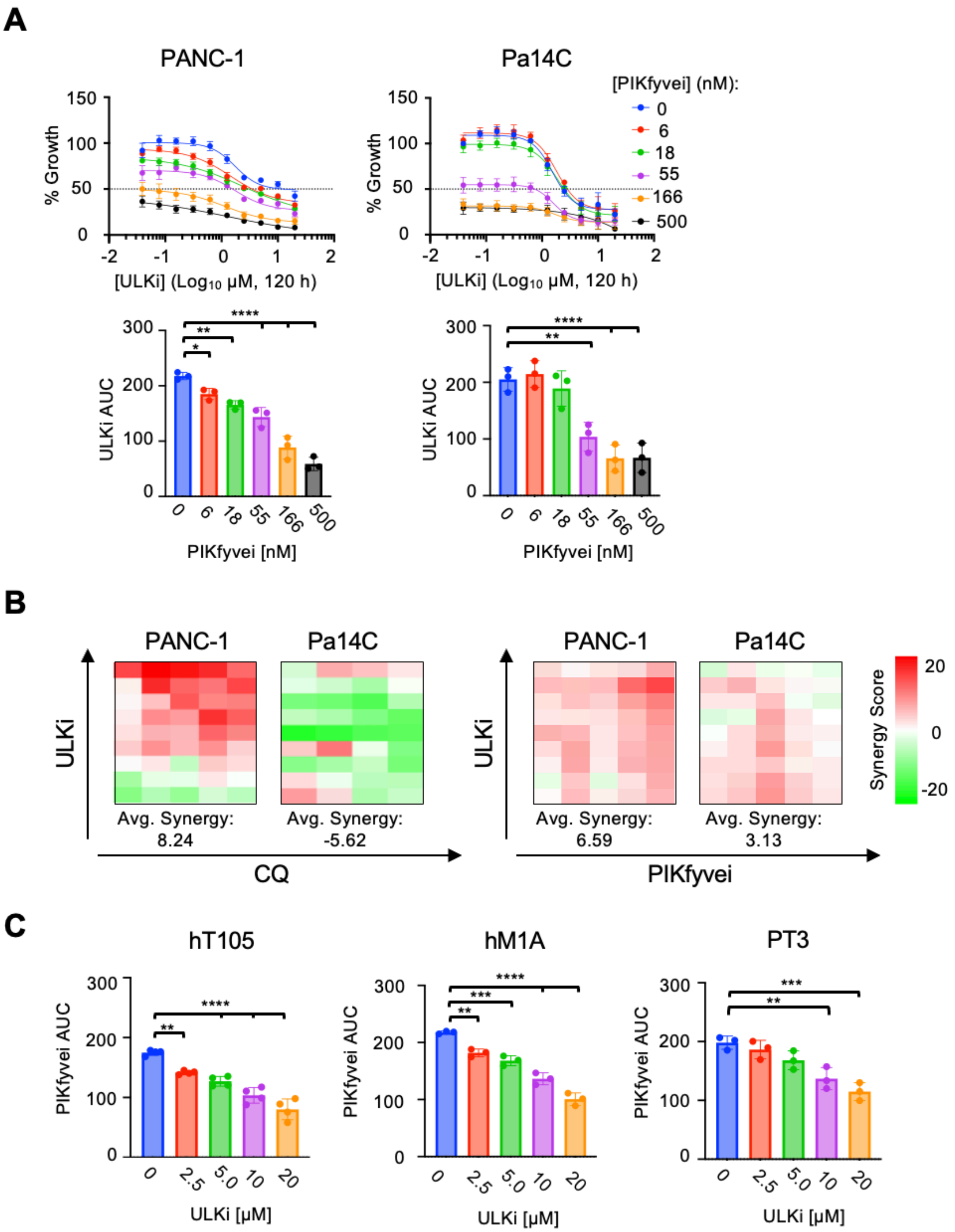

**Supplementary Figure 7**

ULKi sensitizes PDAC cells to terminal stage autophagy inhibition. **A,** Top, five-day viability assay in PANC-1 and Pa14C cells following increasing doses of ULKi (ULK101, [0.078 – 20 μM]) alone or in combination with doses of PIKfyvei (apilimod, [6 – 500 nM]) treatment. Each data point represents mean ± SEM of three independent experiments. Bottom, area under the curve (AUC) values from each biological replicate of data plotted above, error bars denote mean ± SD. Statistical significance was determined by a one-way ANOVA with Dunnett's test. *, $\rho < 0.05$; **, $\rho < 0.01$; ****, $\rho < 0.0001$. **B,** Excess over bliss synergy scores calculated from biological replicate of data in **S5C** and **S7A** using SynergyFinder. Tiles represent synergy score for individual combination. **C,** Area under the curve (AUC) values from each biological replicate of data plotted in **3H**, error bars denote mean ± SD. Statistical significance was determined by a one-way ANOVA with Dunnett's test. *, $\rho < 0.05$; **, $\rho < 0.01$; ***, $\rho < 0.001$; ****, $\rho < 0.0001$.

### Supplementary Fig. S8

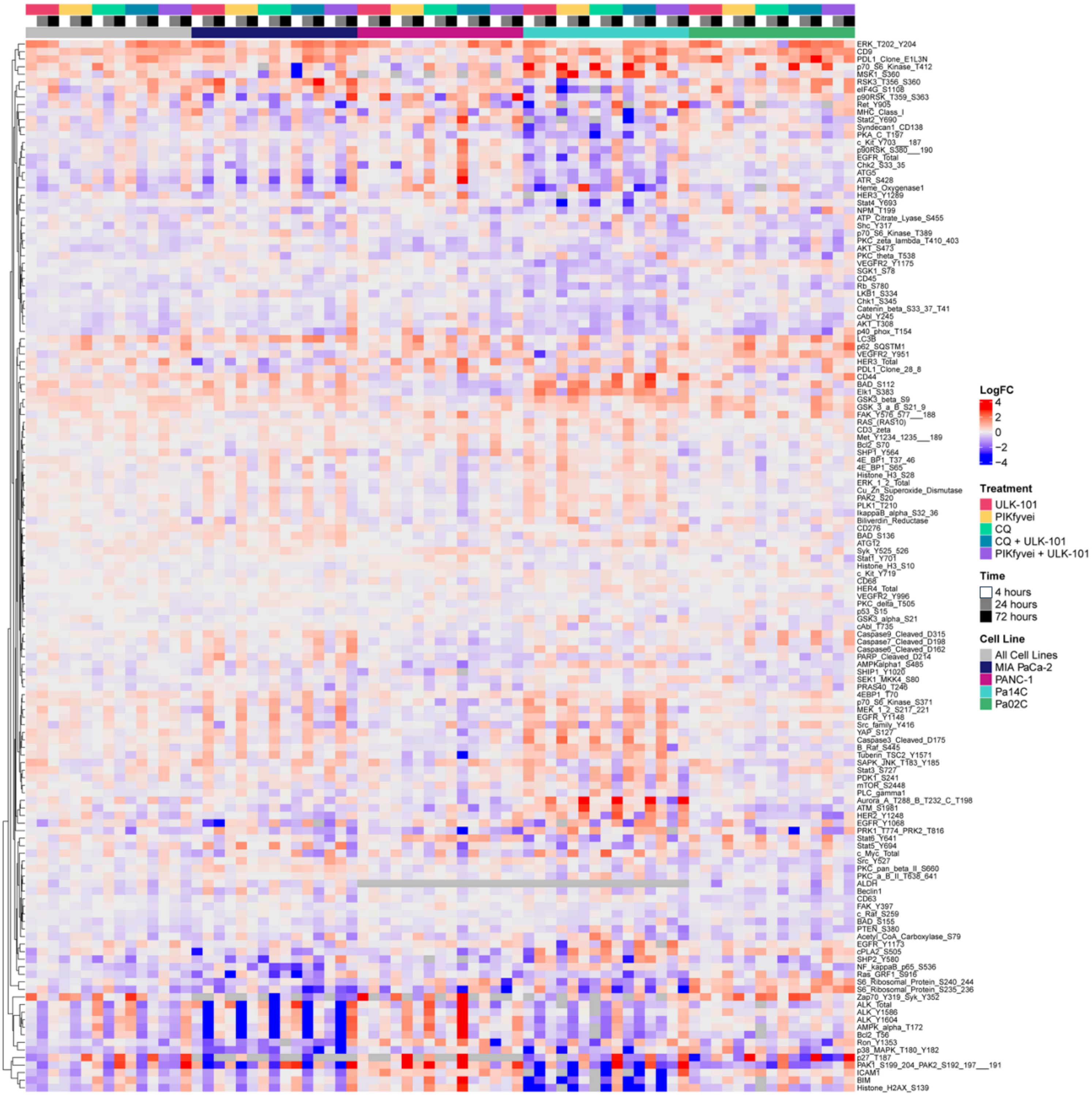

**Supplementary Figure 8**

Autophagy inhibition results in broad signaling changes across PDAC cell lines. Heat map showing median log2(FC) of the 139 proteins/phosphoproteins evaluated by RPPA in Pa14C, Pa02C, PANC-1 and MIA PaCa-2 cell lines treated for 4, 24, 72 hours with vehicle (DMSO), ULKi (3 μM), CQ (6.125 μM), PIKfyvei (apilimod, 100 nM), ULKi (ULK101) +CQ, or ULKi (ULK101) + PIKfyvei (apilimod). The median of four biological replicates for each drug treatment condition is represented, and log2(FC) was calculated compared to DMSO control for each time point.

#### Supplementary Fig. S9

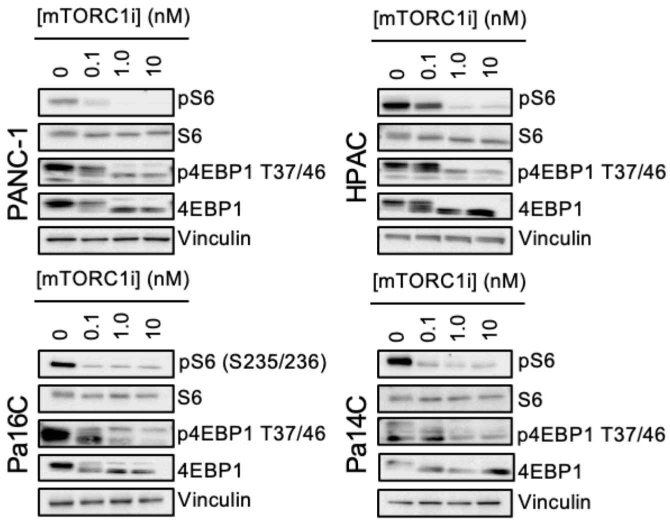

**Supplementary Figure 9**

mTORC1 inhibition with RMC-5552 results in decreased phosphorylation of mTORC1 substrates. Immunoblot of PANC-1, Pa16C, HPAC and Pa14C cell lines following 24 hours of mTORC1i (RMC-5552) treatment at indicated doses. Blots are representative of three independent experiments.

#### Supplementary Fig. S10

**A**

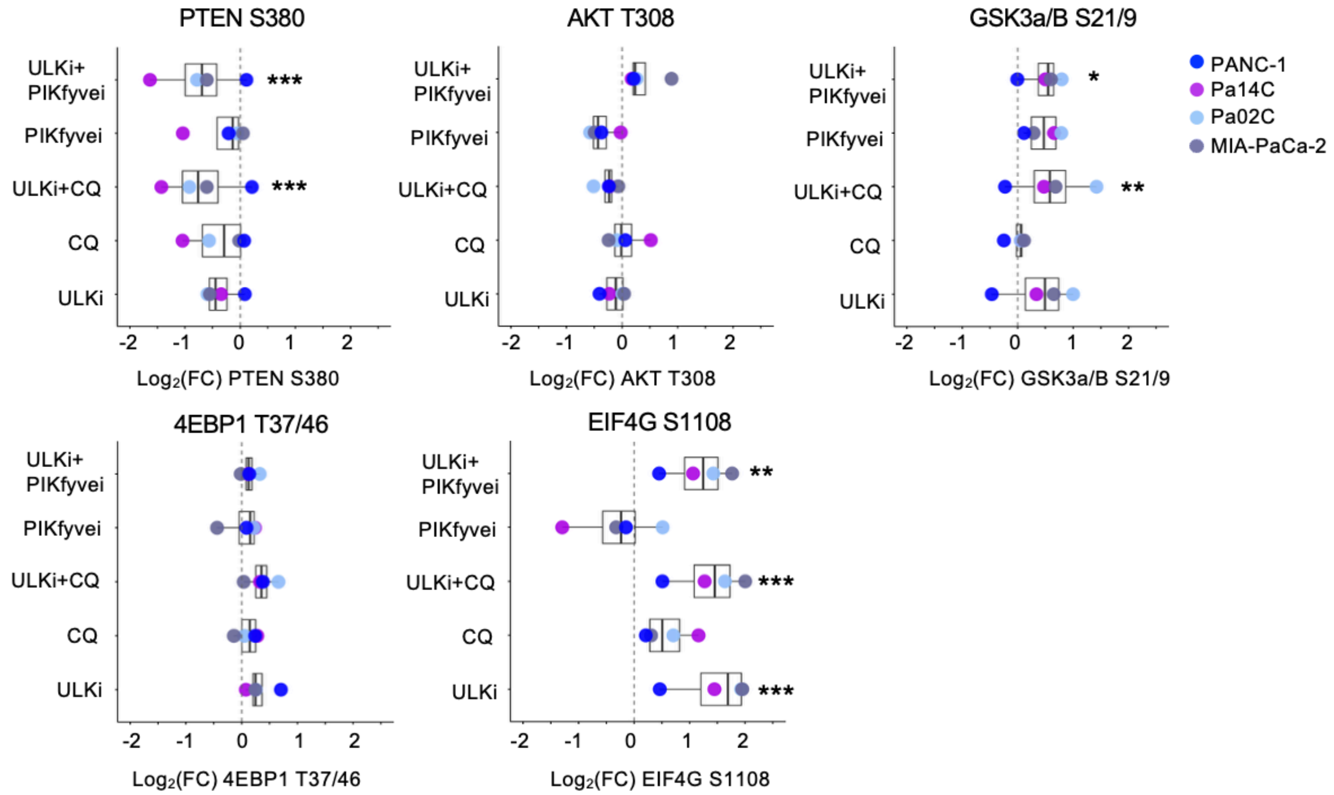


**B**

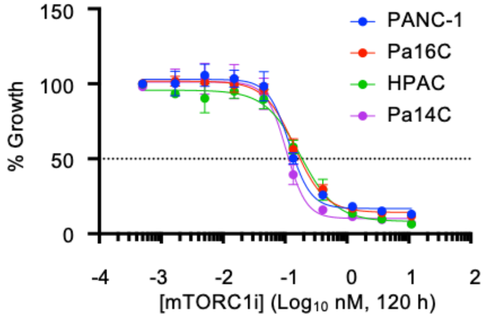


| | PANC-1 | Pa16C | HPAC | Pa14C |
|---|---|---|---|---|
| GI50 (nM) | 0.1414 | 0.1751 | 0.1877 | 0.1145 |

**Supplementary Figure 10**

Vertical autophagy inhibition results in altered activation of PI3K-AKT-mTORC1 signaling pathway components and increases sensitivity to mTORC1 inhibition. **A,** Log2FC values of indicated antibodies from RPPA experiment described in **Fig. 4A** compared to vehicle at 72 hours of treatment. Significance values calculated across all cell lines. *, $\rho < 0.05$; **,$\rho < 0.01$; ***, $\rho < 0.001$. **B,** Five-day viability assay in PANC-1, Pa16C, HPAC, and Pa14C cells following increasing doses of mTORC1i (RMC-5552) Each data point represents mean ± SEM of three independent experiments.

#### Supplementary Fig. S11

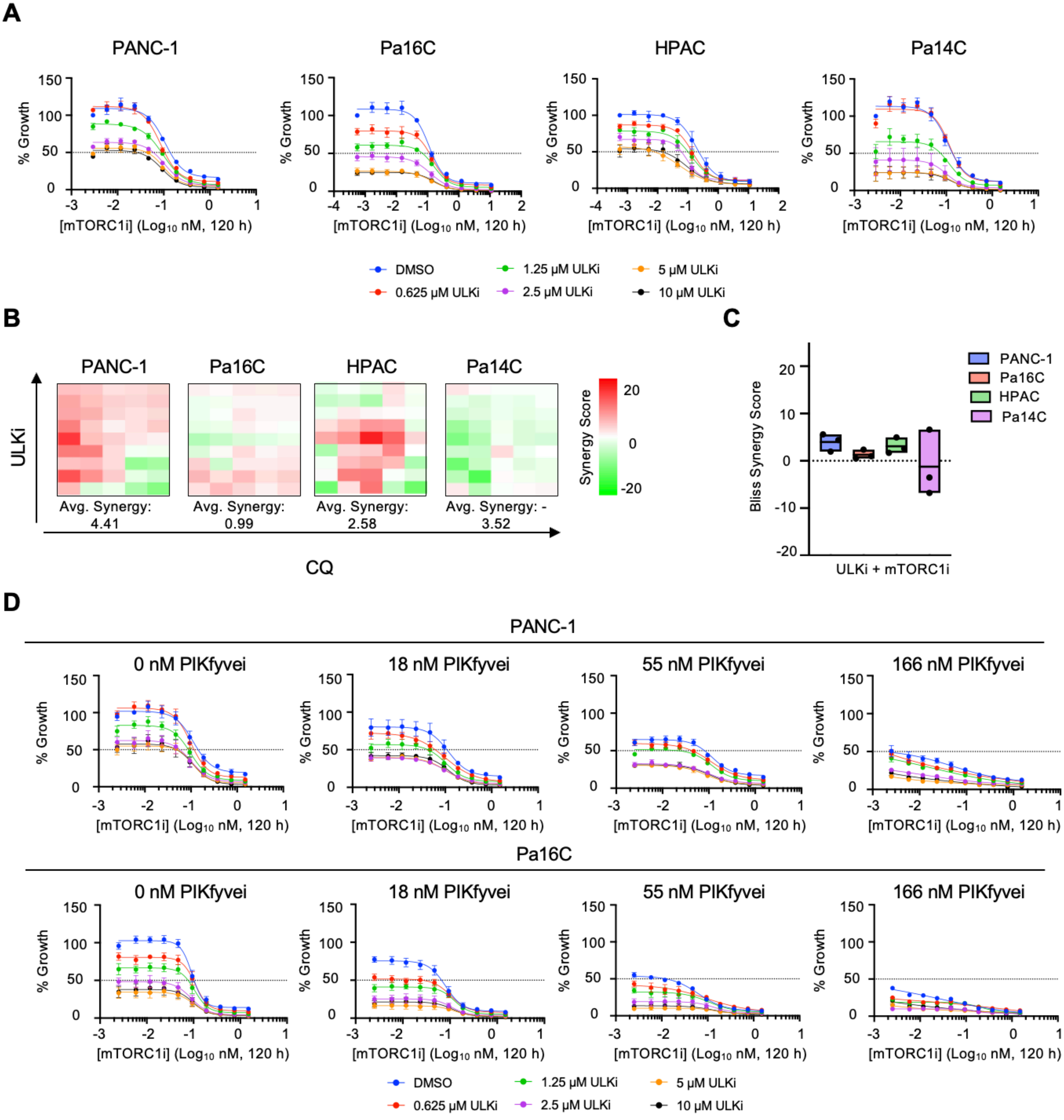

**Supplementary Figure 11**

ULK inhibition synergizes with mTORC1 inhibition to reduce PDAC proliferation and growth suppression is potentiated via PIKfyve inhibition. **A,** Five-day viability assay in PANC-1, Pa16C, HPAC and Pa14C cells following increasing doses of ULKi (ULK101, [0.039 – 10 μM]) alone or in combination with doses of mTORC1i (RMC-5552, [0.005-1.5 nM]). Each data point represents mean ± SEM of three independent experiments. **B,** Excess over bliss synergy scores calculated from biological replicate of data in **A** using SynergyFinder. Tiles represent synergy score for individual combination. **C,** Average synergy scores taken from studies in **A**. **D,** Five-day viability assay in Pa16C and PANC-1 cells following increasing doses of mTORC1i (RMC-5552, [0.005-1.5 nM]) alone or in combination with doses of ULKi (ULK-101, [0.625-10 μM]) treatment, with indicated dose of PIKfyvei (apilimod). Each data point represents mean ± SEM of three independent experiments.
